# Plant diversity stabilizes soil temperature

**DOI:** 10.1101/2023.03.13.532451

**Authors:** Yuanyuan Huang, Gideon Stein, Olaf Kolle, Karl Kübler, Ernst-Detlef Schulze, Hui Dong, David Eichenberg, Gerd Gleixner, Anke Hildebrandt, Markus Lange, Christiane Roscher, Holger Schielzeth, Bernhard Schmid, Alexandra Weigelt, Wolfgang W. Weisser, Maha Shadaydeh, Joachim Denzler, Anne Ebeling, Nico Eisenhauer

**Affiliations:** German Centre of Integrative Biodiversity Research (iDiv) Halle-Jena-Leipzig, Puschstraße 4, 04103 Leipzig, Germany; Institute of Biology, Experimental Interaction Ecology, Leipzig University, Puschstraße 4, 04103 Leipzig, Germany; Computer Vision Group, Faculty of Mathematics and Computer Science, Friedrich Schiller University Jena, Ernst-Abbe-Platz 1-3, 07743 Jena, Germany; Max Planck Institute for Biogeoscience, POB 100161, 07701 Jena, Germany; Institute of Ecology and Evolution, University of Jena, Dornburger Straße 159, 07743 Jena German; German National Centre for Biodiversity Monitoring, Alte Messe 6, 04103 Leipzig, Germany; Department of Computational Hydrosystem, Helmholtz Centre for Environmental Research – UFZ, Leipzig, Germany; Friedrich Schiller University Jena, Institute of Geoscience, Terrestrial Ecohydrology, Burgweg 11, 07745 Jena, Germany; Department of Physiological Diversity, Helmholtz Centre for Environmental Research-UFZ, Permoserstrasse 15, 04318 Leipzig, Germany; Department of Geography, Remote Sensing Laboratories, University of Zurich, Winterthurerstrasse 190, 8057 Zurich, Switzerland; Systematic Botany and Functional Biodiversity, Institute of Biology, Leipzig University, Johannisallee 21–23, 04103 Leipzig, Germany; Terrestrial Ecology Research Group, School of Life Sciences, Department of Ecology and Ecosystem Management, Technical University of Munich, Hans-Carl-von-Carlowitz-Platz 2, D-85354 Freising, Germany

## Abstract

Extreme weather events are occurring more frequently, and research has shown that plant diversity can help mitigate impacts of climate change by increasing plant productivity and ecosystem stability^1,2^. Although soil temperature and its stability are key determinants of essential ecosystem processes related to water and nutrient uptake^3^ as well as soil respiration and microbial activity^4^, no study has yet investigated whether plant diversity can buffer soil temperature fluctuations. Using 18 years of a continuous dataset with a resolution of 1 minute (∼795,312,000 individual measurements) from a large-scale grassland biodiversity experiment, we show that plant diversity buffers soil temperature throughout the year. Plant diversity helped to prevent soil heating in hot weather, and cooling in cold weather. Moreover, this effect of plant diversity increased over the 18-year observation period with the aging of experimental communities and was even stronger under extreme conditions, i.e., on hot days or in dry years. Using structural equation modelling, we found that plant diversity stabilized soil temperature by increasing soil organic carbon concentrations and, to a lesser extent, by increasing the plant leaf area index. We suggest that the diversity-induced stabilization of soil temperature may help to mitigate the negative effects of extreme climatic events such as soil carbon release, thus slow global warming.

---

Extreme weather events are becoming more intense, more frequent, and lasting longer than previously observed^5^. Global climate change has led to changes in soil temperatures and has caused greater variance through climate extremes^6^. Soil temperature affects many physical, chemical, and biological processes and reactions, including water and nutrient uptake^3^, microbial activities, root growth^7^, carbon dioxide flux^8^, ant activity^9^ and plant pests development^10^, thereby affecting seed germination, plant growth and productivity^4^. Fluctuations in soil temperature, including sudden chilling, freezing, or warming, can have dramatic impacts on plants, microorganisms, and soil animals^11^. Thus, mitigating the effects of extreme weather events on soil temperature fluctuations can contribute to stable ecosystem functioning. A few recent studies have shown that plants can buffer air temperature inside forests^12–15^. However, whether plants can contribute to buffering soil temperature is still unclear.

Biodiversity, especially plant diversity, has been shown to enhance ecosystem stability to combat climate change^1^. The biodiversity increases stability hypothesis has been confirmed for several ecosystem functions, including primary productivity^2,16^, the abundance of invertebrates^17^, and trace gas and matter fluxes^18^. However, these studies focused primarily on aboveground processes and rarely investigated soil conditions. Additionally, previous studies on plant diversity and soil interactions focused on the role of soil organisms^19^ and soil nutrients^18^. Little attention has been paid to the effects of plant diversity on soil microclimate^18^, including soil temperature stability. The question of whether plant diversity can reduce soil temperature fluctuation in response to extreme weather and climatic events is of interest because soil temperature regulates many other ecosystem processes, such as soil respiration^20^. Some studies have shown that high plant diversity increases canopy shading^21^ and lowers surface temperature^22,23^ and soil temperature during the growing season^24^. However, there is no study on the effects of plant diversity on soil temperature covering longer continuous time spans. Whether plant diversity plays a role in soil temperature during colder seasons remains largely unexplored. In Central Europe, the consideration of these cold periods is of particular interest, because decomposition processes occur during this time.

Here we report the effects of plant diversity on soil temperature from 2004 to 2021 in a large-scale grassland biodiversity experiment^25^ (the Jena Experiment; see Methods). There has been a large climate variability over these 18 years (Extended Data Figs. 1, 2, Extended Data Table 1). The experimental site contains 84 plots with plant species richness ranging from 1 to 2, 4, 8, 16, and 60, as well as plots with bare soil^25^. Soil temperature was measured automatically at 5 cm and 15 cm depth in each plot with a resolution of 1 minute (Methods), which we convert to a 30-minute resolution for our analysis. This long-term time series allowed us to examine the buffering effects of plant diversity on soil temperature fluctuations within and between days, seasons and years. Here, we investigated two aspects of soil buffering at different temporal scales: (1) soil temperature offset between vegetated and non-vegetated plots at individual time points (Fig. 1); (2) the daily or annual variation/stability of soil temperature (Fig. 3).

**Figure 1.**
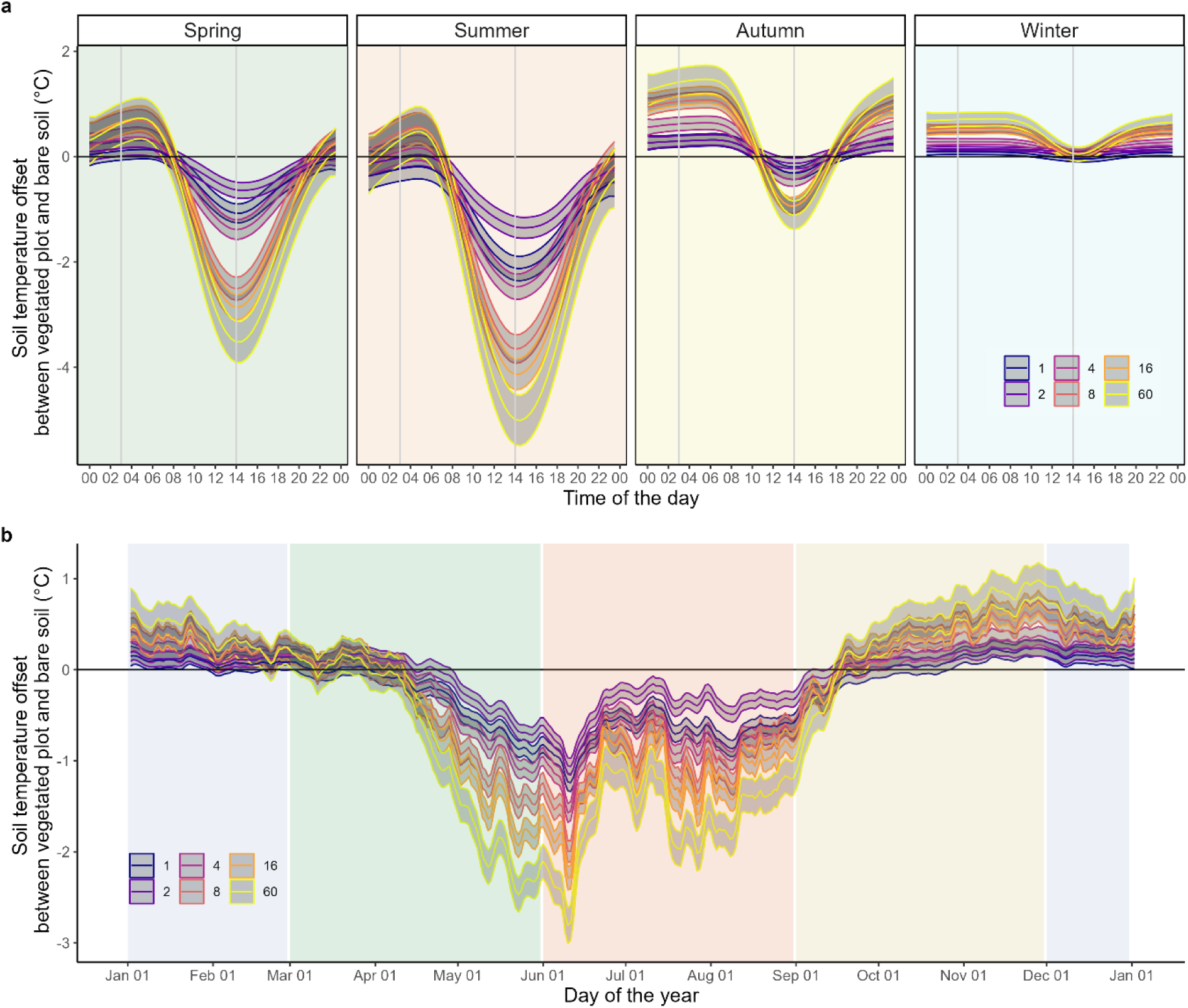
Soil temperature offset between vegetated plots and bare soil at different plant diversity levels (1, 2, 4, 8, 16, and 60 species) on the 30-minute scale within a day for each season (a) and on the daily scale within a year (b). Data with soil temperature at 5 cm depth was shown here. Solid lines and grey shading represent the fitted values and credibility intervals (95%, see Methods). a, Data with a resolution of 30 minutes were used. Annual, monthly, and daily variations were averaged, leaving variations from 80 plots, 48 times per day, and 4 seasons (n = 15,360). Time is Central European Time (CET). b, Daily resolution data were used. Annual variations were averaged, leaving variations of 80 plots and 366 days (n = 29,280).

**Figure 2.**
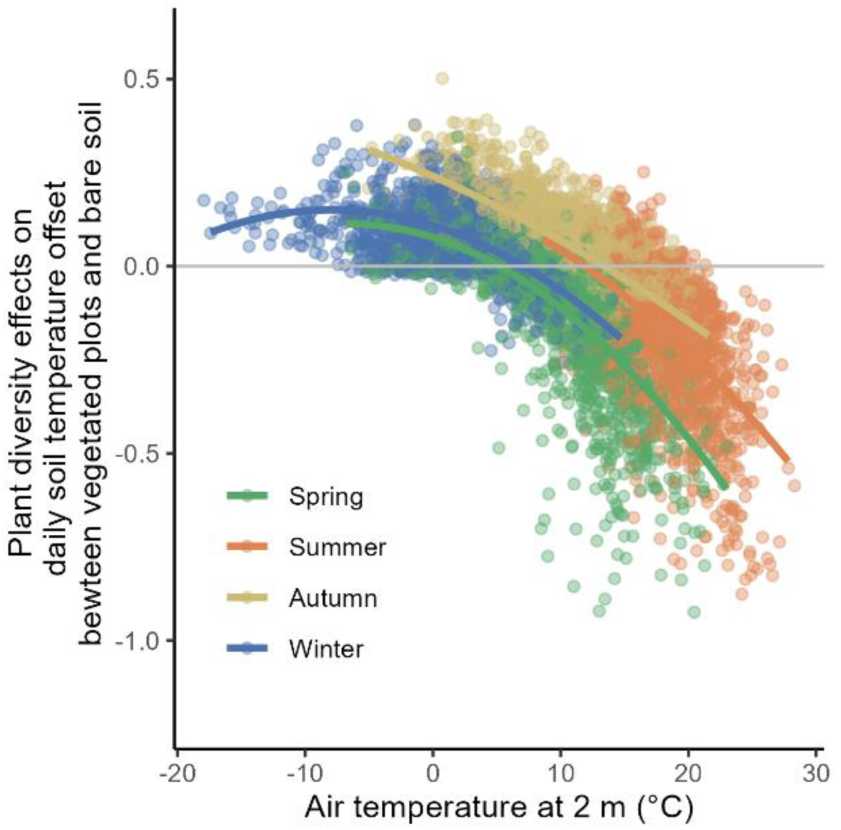
Plant diversity effects on daily soil temperature offset between vegetated plots and bare soil change with air temperature (n = 6,575).

**Figure 3.**
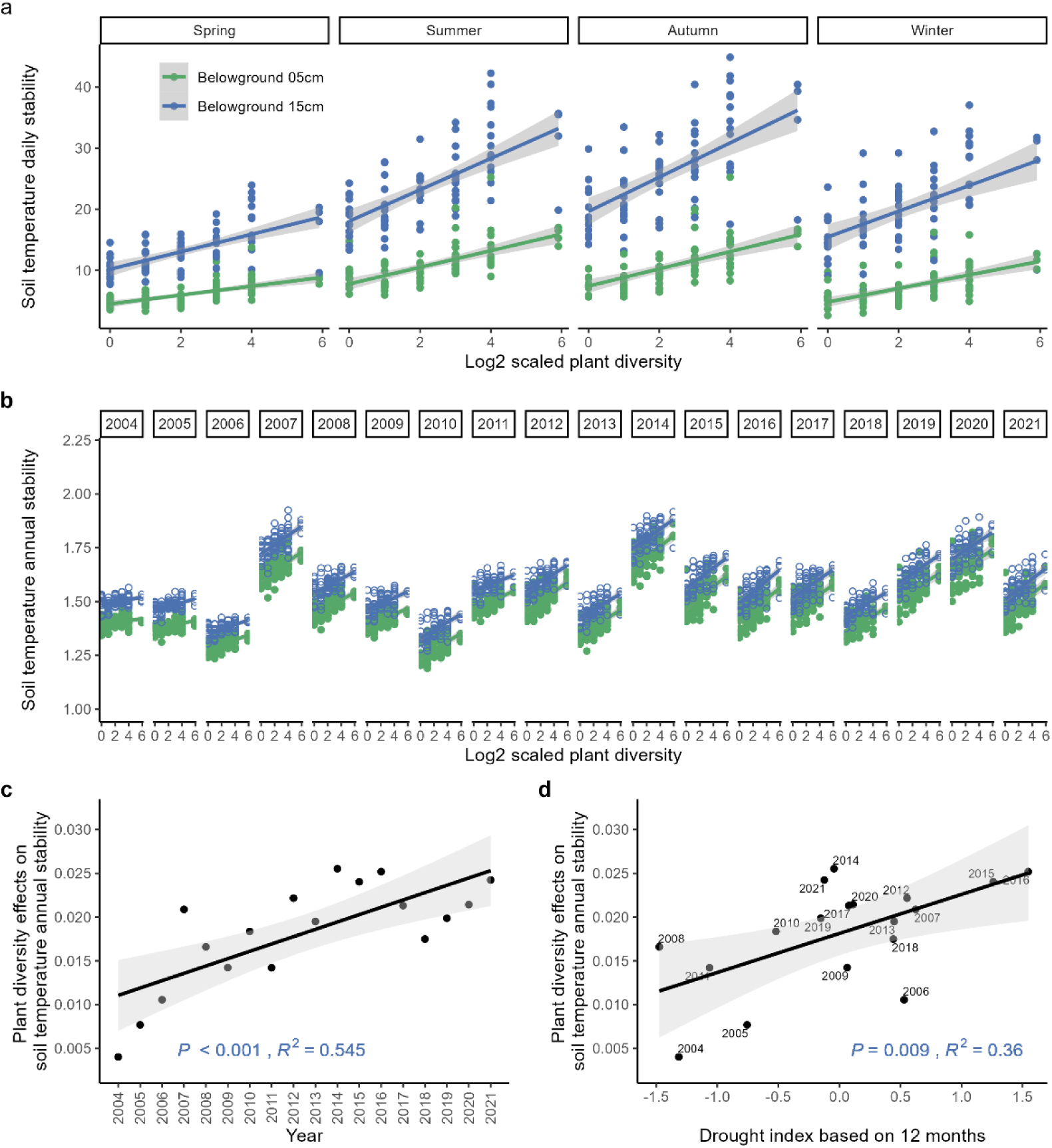
Plant diversity effects on soil temperature stability over the 18 years of the experiment. **a,** The average daily stability of soil temperatures (n = 320); **b,** The intra-annual stability of daily mean soil temperatures (n = 1440). The green lines and the blue lines in **a** and **b** indicate the results at a soil depth of 5 cm and 15 cm, respectively. The plant diversity effect on soil temperature annual stability at 5 cm increased with time since the establishment of the experiment (**c**) and increased with increasing drought (more negative SPEI values) (**d**). The drought index here is calculated by multiplying the SPEI by -1, i.e. the drought situation becomes more severe with increasing values.

First, we explored within-day fluctuations in soil temperature using data with a resolution of 30 minutes. The buffering effects of vegetation on soil temperature were calculated by comparing the soil temperature offset between vegetated plots and bare soil (Methods). A Bayesian time series model was used to test whether the effect of plant diversity changes with time (see Methods). The credibility intervals (95% CI) of the fitted values for the different levels of plant diversity did not overlap (Fig. 1). The higher the diversity of plant communities, the stronger their cooling effect on soil temperature from 12:00 to 16:00 (Central European Time) in spring, summer, and autumn and their warming effect at night (from 02:00 to 06:30) in autumn and winter (Fig. 1a). In summer, when air temperature was highest during the day, soil temperature in 60-species plant communities was 5.01°C [95% CI, -5.49 to -4.53°C] lower than bare soil, which is more than twice the difference between monocultures and bare soil (-2.12°C; 95% CI, -2.35 to -1.89°C) (Fig. 1a). In autumn, when air temperature was lowest, soil in the 60-species plant community was 1.47°C [95% CI, 1.20 to 1.74°C] warmer than bare soil, almost five times the difference between the monocultures and bare soil (+0.32°C; 95% CI, 0.20 to 0.44°C). We also used the offset between soil temperature and air temperature as an additional dependent variable, and found similar effects of plant diversity (Extended Data Fig. 3). In the summer afternoon, soil temperature is higher than air temperature in communities with low plant diversity (+1.09°C; 95% CI, 0.80 to 1.39°C). This may be due to the factor that solar radiation is strongest and the soil is dry at this time, and bare soil heats up faster than air. However, in communities with high plant diversity, the soil is still much cooler than the air (-3.23°C; 95% CI, -3.68 to -2.77°C, Extended Data Fig. 3). This demonstrates that plant diversity can help to stabilize soil temperature on a 30-minute time scale, which in turn may help to stabilize other ecosystem functions.

Second, we focused on daily resolution data to explore the seasonal dynamics of the buffering effect of vegetation, which differs by different levels of plant diversity (Fig. 1b, Extended Data Fig. 4). Even though the seasonal pattern differed from year to year, we found consistent effects of plant diversity (Extended Data Fig. 4). Within one year, the number of extreme heat days and frost days decreased with increasing plant diversity (Extended Data Fig. 5). In spring and summer, the average daily temperature decreased with increasing plant diversity, especially from May to August (Fig. 1b), when air temperature was high and aboveground plant biomass peaked^21^. An exception was the 2-species mixtures, which did not lower the soil temperature during the day as much as the monocultures (Fig. 1a, b). In contrast, in the colder seasons, autumn and winter, plant diversity generally increased soil temperature (Fig. 1b). Although mean soil temperatures were similar in autumn and spring, the variance in spring was much greater, and the direction of the effects of plant diversity was opposite (Fig. 1a, b). During the soil warming period, plant diversity helps to prevent sudden soil warming in spring. In contrast, plant diversity helps to buffer soil temperature from rapid cooling in autumn. Thus, changes in air temperature are propagated more slowly into the soil in more diverse plant communities. Although the effect size was much smaller in winter (Fig. 1a, b), it is nonetheless important because even a small difference can imply freezing vs. non-freezing soil conditions^26^.

**Figure 4.**
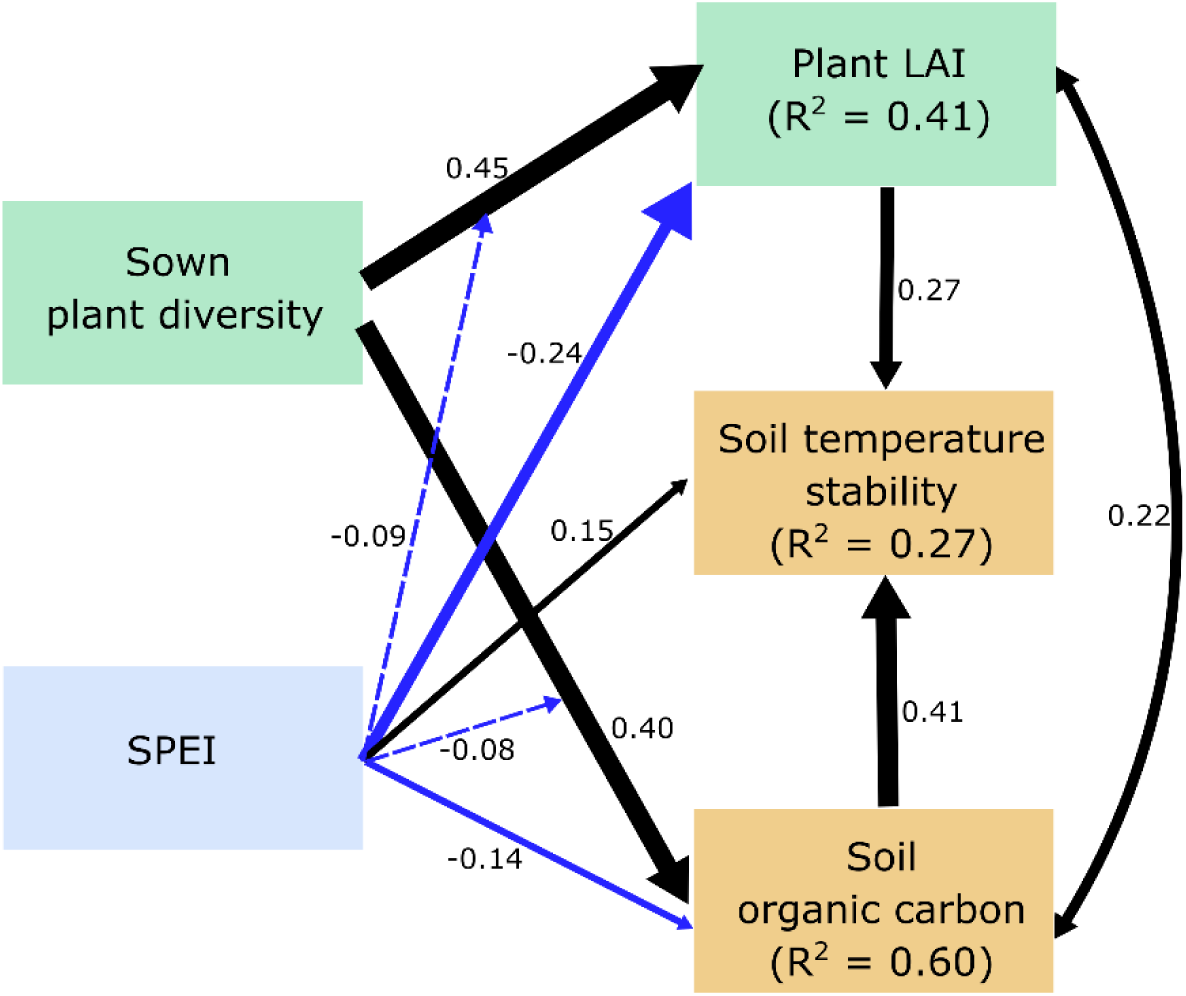
Hypothetical mechanisms underlying significant plant diversity effects on soil temperature stability. A structural equation model (SEM) exploring the effects of plant diversity on intra-annual soil temperature stability across 80 experimental plots through plant leaf area index (LAI) and soil organic carbon (n = 480). Solid black and blue arrows represent positive and negative standardized path coefficients, respectively, and dashed arrows represent interactive effects of plant diversity and drought index. Double-headed arrows indicate covariances. They were included in the model to account for correlations between variables. Standardized path coefficients are given next to each path; widths of significant paths are scaled by standardized path coefficients. In this model, all the paths were significant. Conditional *R*^2^ (based on both fixed and random effects) is reported in the corresponding box. The overall fit of the piecewise SEM was evaluated using Shipley’s test of d-separation: Fisher’s C = 2.768 and *P* value = 0.25 (if *P* > 0.05, then no paths are missing and the model is a good fit).

To calculate the effects of plant diversity on soil temperature offset between vegetated and non-vegetated plots, we fitted a linear regression model at each time point (log-scaled plant diversity as a linear term). We then used the slope of this regression as a proxy for the strength of the effect size (Methods). Plant diversity effects can change rapidly along with changes in meteorological conditions in a short period of time. To test this, we regressed plant diversity effects calculated from daily data on air temperature measured at the climate station at the field site. We found that air temperature (2 m above ground) significantly affected diversity effects (*F_(1,6342)_* = 4304.24, *P* < 0.001, and quadratic term: *F_(1,6342)_* = 698.89, *P* < 0.001, Extended Data Table 2, Fig. 2). The effects of plant diversity were stronger at high air temperatures, suggesting that more diverse communities have a stronger buffering effect on soil temperature at higher air temperatures (Fig. 2). In contrast, on the coldest days, plant diversity effects were not affected by air temperature (Fig. 2). This could be due to snow cover, which helps to insulate soils from cooling at very low air temperature. The interaction between air temperature and season (*F_(3, 6342)_* = 22.36, *P* < 0.001, Extended Data Table 2, Fig. 2) was significant. The negative effects of plant diversity on soil temperatures were strongest in spring and summer (Fig. 2), indicating a direct buffering effect of plant diversity against warmer air temperatures. After accounting for the effects of air temperature and further decomposing the residual variance of the plant diversity effects, we found that the seasons within a year and the hours within a day still explained quite a large part of the variance (Extended Data Fig. 6). This implies that plants not only have an inactive insulating effect that is strongly dependent on the air temperature, e.g. through the vegetation cover, but can also actively regulate the microclimate on an hourly and seasonal level, independent of the air temperature.

While our findings focus mainly on temperature at 5 cm soil depth, we also analysed data collected at 15 cm (all plots) and 60 cm depth (available only in one of the four experimental blocks; Extended Data Figs. 1, 7, 8, 9). Overall, we observed that the effects of plant diversity at deeper soil depths were consistent with the results at 5 cm soil depth, although the effects at 60 cm depth were attenuated (Extended Data Figs. 7, 8), and no longer visible (Extended Data Figs. 9). This result was to be expected, since it is known that deeper soil layers response less immediately to meteorological fluctuations^27^. Given that soil warming has been shown to increase soil carbon loss through enhanced microbial respiration^28,29^, our results suggest that increased plant diversity could buffer soil temperatures from sudden changes at different soil depths in the short term to mitigate the effects of climate change on soil microbial communities and carbon release.

On a longer temporal scale, we analysed the stability of soil temperature. To understand the effects of plant diversity on within-day and between-day within-year soil temperature stability, we calculated daily and intra-annual soil temperature stability for an accumulated period by dividing the mean soil temperature by its standard deviation (*^µ/^σ*) derived from the 30-minute and daily mean soil temperature data, respectively (Methods). The main effect of plant diversity was significantly positive, i.e., plant diversity significantly increased soil temperature stability at both soil depths (i.e., 5 and 15 cm; Fig. 3a, b; F_(1,75)_ = 89.39, *P* < 0.001 at daily time scale; F(_1,75)_ = 105.81, *P* < 0.001 at annual time scale), indicating a constant buffering effect of plant diversity throughout the day and year. At annual scale, there was no significant interaction between plant diversity and soil depth (Fig. 3b, F_(1,78)_ = 0.015, *P* = 0.90), highlighting the consistency of plant diversity effects. However, at the daily scale, the effects of plant diversity are stronger at a soil depth of 15 cm (Fig. 3a, F_(1,78)_ = 9.29, *P* = 0.003). This means that plant diversity also affects the soil layer from 5 to 15 cm, which further reduces the soil heat flux and stabilizes the soil temperature.

We also found that the positive effects of plant diversity on soil temperature intra-annual stability became more substantial with time after the establishment of the experiment (Fig. 3c, F(_1,15)_ = 23.81, *P* < 0.001), which is consistent with the analysis of daily soil temperature offset (F_(1, 16)_ = 24.57, *P* < 0.001, Extended Data Table 2, Extended Data Fig. 8). This is also in line with the increasing plant diversity effects on plant productivity observed in many ecosystems^30–32^. This result supports that biodiversity effects increase over time, which implies a high value of old grasslands with a high diversity of plant species.

In addition to a linear trend in plant diversity effects over the 18 years of the experiment (linear effect of ‘year’), annual climate showed considerable variation (Extended Data Fig. 2), which also resulted in annual variation in the buffering effect of plant diversity. After statistical consideration of the linear trend, the drought index “standardised precipitation evapotranspiration index” (SPEI)^33^ still explained a significant portion of the variance in the effect of plant diversity (Fig. 3d, F(1,15) = 4.89, *P* = 0.04). This suggests that, even though the effect of plant diversity strengthened over time, the buffering effect of plant diversity was stronger in years with harsher climates (e.g., drought years). In turn, this result confirms that plant diversity–soil temperature stability relationships are climate dependent^34^.

To investigate the underlying mechanisms of plant diversity effects on soil temperature stability, we used above-and below-ground variables to construct a structural equation model (SEM) (Fig. 4, Methods). Overall, plant leaf area index (LAI), soil organic carbon (SOC), and annual standardised precipitation-evapotranspiration index (SPEI) explained 27% of the variation in intra-annual soil temperature stability. Plant diversity significantly increased plant LAI and SOC, which stabilized soil temperature throughout the year. The direct effect of plant diversity on soil temperature stability was not significant (not included in the SEM, *P* = 0.25), suggesting that most of the plant diversity effect was mediated indirectly through plant diversity-enhanced LAI and SOC. The standardized indirect effect of plant diversity by SOC (0.41) was even higher than that by LAI (0.27). This suggests a strong thermal mediation of SOC to stabilize the belowground environment against climate fluctuations and thus possibly against longer-term climate change and variability. SOC has been shown to be related to increased soil porosity^35^. Higher soil porosity can improve thermal diffusivity, an indicator of the rate at which a change in temperature is transmitted through the soil by heat conduction^36^. Thus, the higher the SOC, the slower the temperature change is transmitted to deeper soil layers^35^. In the Jena Experiment, researchers have found that the positive effect of plant diversity on SOC expanded to deeper soil layers^37^. With higher plant diversity, there are more SOC at both 5 cm and 15 cm, thus more insulation effects at 15 cm, which explains the stronger effects in the deeper soil layer of 15 cm than 5 cm (Fig. 3a). LAI is an important indicator of canopy structure^23^, which affects the insulating effect. Plant diversity increases LAI and plant communities of higher LAI help to reduce solar radiation, increase albedo and affect wind speed, which in turn reduces heat fluxes^23^. LAI is also highly correlated with plant productivity^25^, which is associated with an active cooling effect in hot weather, e.g., through evapotranspiration^13^. Taken together, these results provide evidence that plant diversity enhances soil temperature stability by increasing both the aboveground plant leaf area and SOC. This SEM model also shows that climate (drought index SPEI) modulates the effect of plant diversity on LAI and SOC. These interaction effects explain the former result that the effects of plant diversity on soil temperature stability are stronger in drier years (lower SPEI) (Fig. 3d).

In summary, we found the first evidence of a stabilizing effect of plant diversity on soil temperature across temporal scales. Our results show that the effect of plant diversity increased over time after the establishment of the experiment. The magnitude of the effect of plant diversity on soil temperature stability was higher on days with high air temperatures and in dry years than on days with moderate temperatures and in normal years, respectively. These buffering effects of plant diversity on soil temperature reveal a mechanism by which plant diversity can reduce the impacts of extreme weather events on soil temperature and thus protect soils from heat, drought stress and frozen damage. Future climate modelling should incorporate these plant diversity effects on soil to improve the prediction of climate impacts on natural ecosystems. Our results also point to the further potential of using plant diversity as a nature-based solution to climate change mitigation. Because many biological (e.g., microbial or macro-organism activities, plant root growth), chemical (e.g., cation exchange capacity, soil carbon and available nutrients, soil pH), and physical (e.g., soil structure, aggregate stability, soil moisture) processes are strongly dependent on soil temperature and its stability over time^4^, a more stable soil environment may slow potential positive climate feedback effects. This also highlight plant diversity as a crucial ecosystem property that contributes to the continuous provision of multiple ecosystem functions.

## Online content

Methods and additional Extended Data display items are available in the online version of the paper; references unique to these sections appear only in the online paper.

## METHODS

### Study site and experimental design

The Jena Experiment is a large-scale, long-term grassland experiment initiated in spring 2002 and measures several variables across an experimental plant diversity gradient^25^. It is located in the Saale River floodplain near the city of Jena (Thuringia, Germany; 50°55′N, 11°35′E, 130 m a.s.l.)^38^. The mean annual air temperature at the experimental site was 9.8°C, while the mean annual precipitation was 571 mm, calculated from the measurements of the climate station at the Jena Experiment site from 2004 to 2021. The main experiment of the Jena Experiment used a completely randomised block design. It consists of 86 plots, divided into four blocks to account for the different soil conditions^38^. The treatment levels of plant species richness from 0 to 60 were randomly allocated to the plots within each block. Initially, each plot had an area of 20 x 20 m. In 2010, the plot size was reduced to 104.75 m^2^ by terminating subplot treatments (the core area is 6 x 5.5 m)^25^. The Jena Experiment comprises 60 plant species belonging to four functional groups (i.e., grasses, small herbs, tall herbs, and legumes) typical for semi-natural grasslands in the study region. Vegetation plots include a gradient of plant species richness (1, 2, 4, 8, 16, and 60 species). All species richness levels are represented by 16 replicates, except for the 16-species mixtures, which had only 14 replicates (the number of legume and small herb species included was less than 16), and the 60-species mixtures, which had four replicates^38^. Our sensitivity analysis shows that, the results and conclusions do not change significantly even if we excluded 60-species mixtures (Extended Data Fig. 10). Two monoculture plots were abandoned in later years due to poor coverage of target species, which resulted in 80 vegetation plots and an additional four bare ground plots in our analysis. The plots were mowed twice a year, and the harvested plant material was removed. All plots were not fertilized, but weeded regularly (two to three times per year) to maintain the composition of target species.

### Soil temperature and climate data collection

Soil temperature at 5 cm and 15 cm was measured since 2003 with thermometers of the CAN-bus module system (JUMO, Germany). Since plants needed some time to establish themselves, we used the data from 2004 onwards for our analysis. The temperature sensors are lance probes with a diameter of 4.5 mm and a length of 200 mm. The measuring element is a PT100-resistor with a tolerance of 1/3 DIN, which means +/-0.1°C at 0°C. The sensor operates in a 4-wire-connection to the data acquisition module of the CAN-bus network. There is no wrapping around the sensor. In addition, 22 plots in the block II, covering the entire gradient of plant diversity, were equipped as intensively measured plots. Additional sensors were installed^25,38^ to measure the soil temperature at the depth of 60 cm (Extended Data Fig. 1).

Furthermore, a climate station in the centre of the field site records many climate variables, such as soil surface temperature, air temperature, relative humidity at 2 m height, soil water content, precipitation, total downwards radiation, and total upwards radiation (infrared temperature sensors Heitronics KT 15). The data from this climate station show that the climate has changed over these 18 years, as evidenced by a significant increase in air and soil temperature (Extended Data Fig. 2). While the resolution of the soil temperature measurement at plot level is 1 minute, the climate station recorded data every 10 minutes. For our analysis, we converted data to a 30-minute resolution and then calculated the daily mean and variance based on this resolution. For all data, Central European Time (CET) was applied to the temperature measurement. CET is one hour ahead of Coordinated Universal Time (UTC).

### Data pre-processing and quality control

Since our data were collected over ∼18 years (with a total of approximately 129 million individual microclimate measurements per year), we had to account for measurement errors that in rare cases persist over several years. We solved this by applying two distinct filters to the raw data with 1-minute resolution. First, we filtered values in an unreasonable range (e.g., temperatures above 50°C) with a simple threshold. Second, we calculate the whiskers of a boxplot (1.5 IQR) for each minute in our data to identify and filter out outlier plots that are anomalous based on the temperature and the variance of all other plots. With this 1-minute resolution dataset, the 30-minute resolution could be derived by averaging while excluding missing values. The daily resolution dataset was then derived from this 30-minute resolution dataset.

While data gaps do not affect the 30-minute dataset, the missing data must be filled in for the daily and annual analysis so that the dataset is not biased due to large gaps (e.g., the annual temperature could be unreasonably high if many winter measurements are missing). To achieve this, we calculated the mean of all available plots in this specific 30-minute interval in the same year and use it as a filling value. However, some gaps (8%, Extended Data Table 1, Extended Data Fig. 1) extended over all plots (e.g., due to a flood in 2013). For these gaps, we calculate the mean of all the plots during other years and use these values to fill them. It is important to note that both the cleaning and filling methods are conservative, as they do not distinguish between levels of plant diversity. This means that our approach reduces the difference between the different levels of plant diversity. We also performed sensitivity analyses by excluding the years in which more than 15% of the values were missing. The results and conclusions from these analyses do not change (Extended Data Fig. 11).

### Derived data calculation

With 30-minuite resolution data, we calculated the buffering effect of vegetation by subtracting the mean soil temperature of the four bare soil plots from the soil temperature of each vegetation plot for each time point, which leaves us with the soil temperature offset between the vegetation plot and the bare soil (Fig. 1a). We also calculated the temperature offset between the soil temperature and the air temperature, using the air temperature as a reference (Extended Data Fig. 3).

Then we aggregated the data to daily level (Fig. 1b, Extended Data Fig. 4) and fit a linear regression to the relation between the daily mean soil temperature offset and the log-scaled plant diversity (predictor variable). The slope of this regression is used as a proxy for the plant diversity effect on buffering soil temperature. These approximations are then plotted against air temperature on a given day (Fig. 2, Extended Data Fig. 7) and against time (Extended Data Fig. 8).

For both daily and annual soil temperature buffering effects, we used a dimensionless measure of ecosystem stability, quantified as the ratio between the mean and standard deviation (µ/σ) of soil temperature over hours within a day, or over days within a year.

### The standardised precipitation evapotranspiration index (SPEI)

For our analysis of drought impacts on the annual buffering effects of plant diversity, we used the SPEI^33^ to compress drought severity into a single variable^1^. The SPEI is a well-established drought index that includes precipitation, temperature, and evapotranspiration. To use the most accurate estimate, we calculated it manually based on data from the local climate station at the field site of the Jena Experiment. For this calculation, a time series of the climatic water balance (precipitation minus potential evapotranspiration) is required. The monthly mean/maximum/minimum air temperature, incoming solar radiation, saturation water pressure, atmospheric surface pressure, and precipitation were used to estimate the reference evapotranspiration (ET0), which is considered equivalent to potential evapotranspiration (PET). PET is the amount of evaporation and transpiration that would occur if a sufficient water source were available. We calculated the ET0 with the “penman” function in the “SPEI” package in R^39^, which calculates the monthly ET0 according to the FAO-56 Penman‒Monteith equation described in Allen et al. (1998)^40^. We considered annual water balances and thus used SPEI-12^1,33^, which was calculated on an annual time scale, for our annual analysis of soil temperature stability (Extended Data Fig. 2d).

### Biotic and abiotic covariate data

In addition, data of variables such as plant aboveground biomass, plant cover, leaf area index (LAI), root biomass, soil organic carbon (SOC), microbial biomass, and soil basal respiration were collected for further analysis to investigate the underlying mechanism of the plant diversity – soil temperature stability relationship. Plant aboveground biomass, plant cover, and LAI are highly correlated^23^. The precision of the plant cover data is limited, as it is only estimated as a percentage of the total vegetation area by eye. Since plant aboveground biomass could not reflect the distribution of leaf area and canopy vertical structure in the plot, we chose LAI to represent the aboveground leaf area coverage.

#### Plant LAI

was measured in August, corresponding to peak aboveground plant biomass. LAI was measured before mowing in the central area of the plot using a LAI-2000 plant canopy analyser (LI-COR Inc., Lincoln, Nebraska, USA) by taking one reference measurement above the canopy and ten measurements approximately at 2 cm above the ground along transects^41^.

#### Soil water content

was measured by frequency domain reflectometry (FDR) using a portable FDR profile probe (PR1/6 and PR2/6, Delta-T Devices Ltd., Cambridge, UK)^42^. Measurements were taken at approximately weekly intervals during the growth season (April–September) and biweekly in other months from 2004 to 2021 with an interruption in 2006, 2007, and 2019.

#### Soil microbial biomass carbon

was determined from 2004 to 2021, except 2005^43^, using an O2-microcompensation apparatus^44^. Soil sample of approximately 5 g of soil (fresh weight) in each plot were collected in May each year.

#### Standing root biomass

was sampled in June 2003, 2004, 2006, 2011, 2014, 2017, and 2021. At least three soil cores were taken per plot in each year, and soil cores in each soil layers were pooled plot-wise. We only used the root biomass at the soil depth of 0 – 5 cm in the SEM. Roots were washed, dried, weighted, and calculated as grams of dry mass per square metre. For details, please see Ravenek et al., 2014^31^.

#### SOC

was measured in April 2003, 2004, 2006, 2011, 2014, and 2017. Three soil samples (4.8 cm in diameter, 0–30 cm deep) were taken per plot using a split-tube sampler (Eijkelkamp Agrisearch Equipment, Giesbeek, The Netherlands)^45^. In our SEM analysis for soil temperature at 5 cm, only 0-5 cm SOC was used. The soil was dried, sieved (2 mm mesh), and milled. The total carbon of the soil samples was determined by an elemental analyser after combustion at 1,150°C (Elementar Analysator vario Max CN, Elementar Analysensysteme GmbH, Hanau, Germany). Inorganic carbon concentration was measured by elemental analysis after removing organic carbon by oxidation in a muffle furnace at 450°C for 16 h. The organic carbon concentration was calculated from the difference between the total and inorganic carbon concentrations.

### Statistical analyses

Time series analysis was performed using R-INLA (R-Integrated Nested Laplace Approximation)^46^. To compare the effects of plant diversity over time, we modelled the soil temperature offset between vegetated plot and bare soil as a function of plant diversity effects and a trend over time. The model is given below.

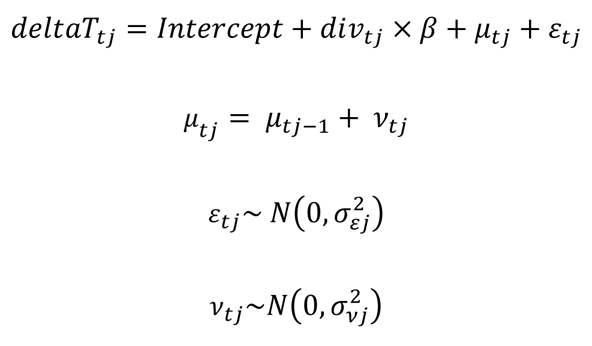

The *deltaT_t_* is the soil temperature offset between the vegetated plot and the bare soil at time t. The *div_t_* is the categorical variable plant diversity, which has six levels. It allows for a different mean temperature offset per plant diversity level. The trend *µ_tj_* is modelled as a rw1 random walk trend based on a penalized complexity prior^46^ with parameters of U = 1 and α = 0.01. Here, we allowed separate trends for each plant diversity level j to investigate whether the trends differ with plant diversity.

We have two time series datasets for this time series analysis. One is the 30-minute intraday resolution data for each season (n = 48 × 4 × 80) to observe the daily pattern (Fig. 1a). The other is the daily data, averaged over the 18 years (n = 366/365 × 80, Fig. 1b).

For 18 years of daily data, we analysed the effects of plant diversity as a function of air temperature using mixed models and summarised results in analyses of variance (ANOVA) tables (Extended Data Table 2). The fixed terms in the model were the air temperature from the climate station [linear (Tair) and quadratic contrast (qTair)], the season (factor with four levels), the centralised linear year (cyear), and interactions of these terms. The random terms were the year, the months within year, and the autocorrelation of the plant diversity effects between days within each year.

After accounting for the effects of air temperature, we explained the residual variance in the effects of plant diversity by different temporal variables, i.e. year, seasons within a year, months within a season, days within a month and hours within a day. We used sequential (type I) sums of squares and calculated the proportion of the total sum of squares that each temporal variable explained.

A linear mixed-effects model was built to test the effects of the logarithm of plant diversity and soil depth on soil temperature stability. For annual soil temperature stability, the block was fitted as a covariate first to exclude the variation of the random position, then the logarithm of plant diversity, soil depth, a centralised linear year and their interactions in the fixed term were fitted. The random term is the nested structure of plot and soil depth, as well as the interaction of plot and year. For daily soil temperature stability, the year was replaced by the season.

A simple linear regression was used to study the contribution of time and climate to the effects of plant diversity on annual soil temperature stability. In the fixed term, the centralized linear year was fit first, followed by the drought index (SPEI). Sequential (type I) sums of squares were used, which means that the effects of the drought index (SPEI) were corrected for the linear year.

### Structural Equation Model (SEM)

Since belowground variables soil organic carbon and root biomass were sampled only once in two or three years, we used only the years (2004, 2006, 2008, 2011, 2014, 2017) that contained the belowground information data for the SEM modelling. SEM was designed to investigate the underlying mechanisms of the significant plant diversity effects on soil temperature stability. To formulate hypotheses about pathways in the model, we searched the literature for knowledge on soil temperature stability and conducted mixed-effects modelling to estimate the effects of covariates on soil temperature stability.

Previous studies have shown that thermal diffusivity is an indicator of soil temperature stability, because it indicates the rate at which a temperature change is transmitted by conduction through the soil^36,47^. Temperature changes are transmitted rapidly through the soil when the thermal diffusivity is high. In addition, research shows that higher soil organic carbon content (SOC) increases soil porosity^35^, which reduces soil thermal conductivity and diffusivity, especially when soil pores are filled with air. As a result, SOC acts as an insulator, and the presence of SOC cools the soil in summer and has a warming effect in winter^36^. Similarly, aboveground plant leaf cover can also act as an insulator to stabilize soil temperature^23^.

Initial mixed-effects models modelling the effects of covariate data on annual soil temperature stability were performed in R (Extended Data Fig. 12). It can be seen that only LAI, root biomass, and soil organic carbon have a positive relationship with soil temperature stability. Soil water content has a strong positive effect on the thermal conductivity as well as on the heat capacity. The wetter the soil, the higher the thermal conductivity and heat capacity^47^. Since thermal diffusivity is the ratio of thermal conductivity to volumetric heat capacity, thermal diffusivity can be less sensitive to the soil water content^47^. Therefore, we didn’t include the soil water content in the SEM.

Data from LAI measurements in August were used in the SEM, because peak plant growing season LAI can represent aboveground annual net primary productivity.

Given these preparatory analyses and considerations, we have only included SOC and LAI in August in our final SEM model (Fig. 4). Since we have data from several years, we also included the main effect climate (SPEI) and its interaction with plant diversity in our model. Furthermore, plot was considered as a random factor variable. After optimisation, the statistically non-significant (*P* > 0.05) paths were excluded from the model. Since the chi-square was not significant (*P* > 0.05), we concluded that the model had a good fit. In addition, the conditional R^2^ value was calculated for each general linear mixed-effects model considering both fixed and random terms.

All analyses were conducted using R 4.2.2. The package “INLA” was used for the Bayesian-based time series analysis. The R package “nlme” was used for the mixed-effects models with temporal autocorrelation, while “lme4” and “lmerTest” were used for mixed-effects models with cross random effects. The package ‘piecewiseSEM’ was used for the structural equation model.

## Data and code availability statement

The data and codes supporting the results of this study are deposited in the Jena Experiment Information System (https://jexis.idiv.de/) and will be published after acceptance of the manuscript. The accession codes will then be provided.

## Extended Data are available in the online version of the paper

## Acknowledgements

We thank all the fieldworkers of the Jena Experiment and Dr. Jes Hines for suggestions on data analysis. The Jena Experiment was funded by the Deutsche Forschungsgemeinschaft (DFG, FOR 5000). We gratefully acknowledge the support of iDiv, which is funded by the German Research Foundation (DFG– FZT 118, 202548816).

## Author contributions

E.A., O.K. and K.K. installed and maintained the soil temperature measurement system. N.E. provided the funding and dataset. Y.H. conceived the project; Y.H., G.S. D.H. cleaned and analysed the data. Y.H. and G.S. wrote the first draft of the manuscript. A.E. is the scientific coordinator of the Jena Experiment. G.G., A.H., M.L. C.R. B.S. A.W. W.W. originally created the dataset of the covariate variables. D.E. contributed to time-series analysis with the Bayesian approach. All authors contributed to the development of the ideas, discussed the analysis and results, and edited the manuscript text.

## Ethics declarations

### Competing interest declaration

The authors declare no competing financial interests.

Correspondence and requests for materials should be addressed to Y.H.

## Extended data figures and tables

**Extended Data Figure 1.**
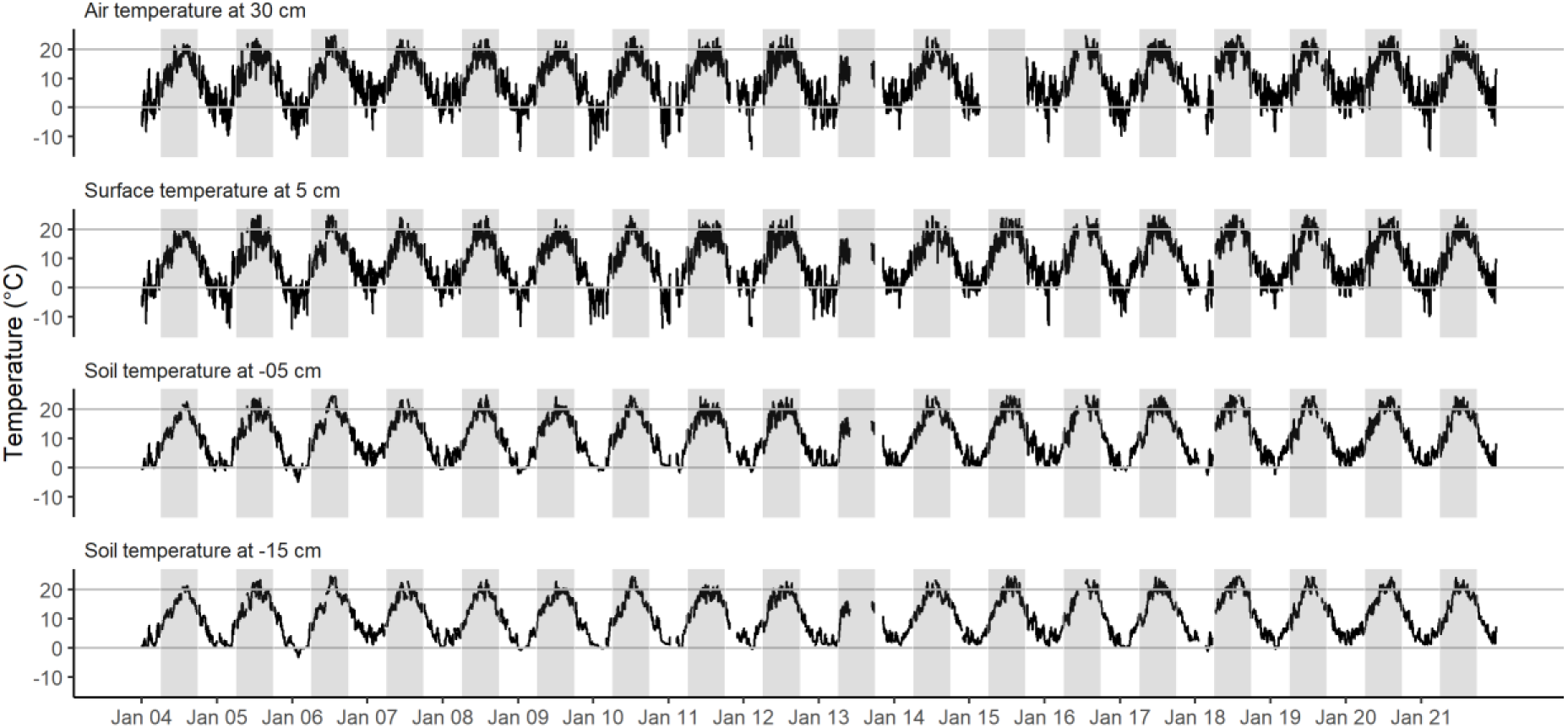
Temperature time series at different heights and soil depths (data from plots in block II).

**Extended Data Figure 2.**
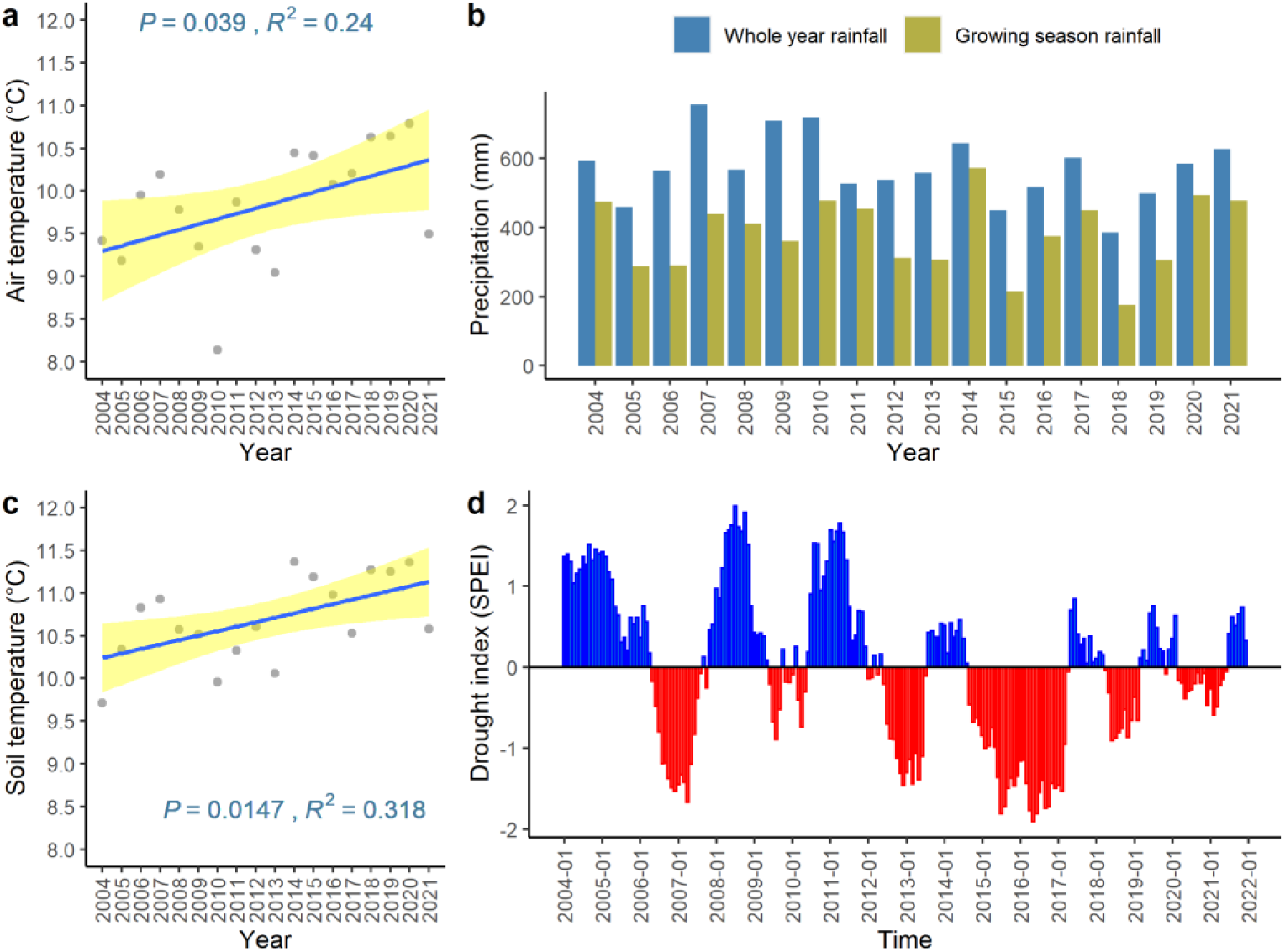
Air temperature at 2 m (**a**), precipitation (**b**), soil temperature at 8 cm (**c**), and drought index (SPEI) (**d**) change with time at the field site of the Jena Experiment.

**Extended Data Figure 3.**
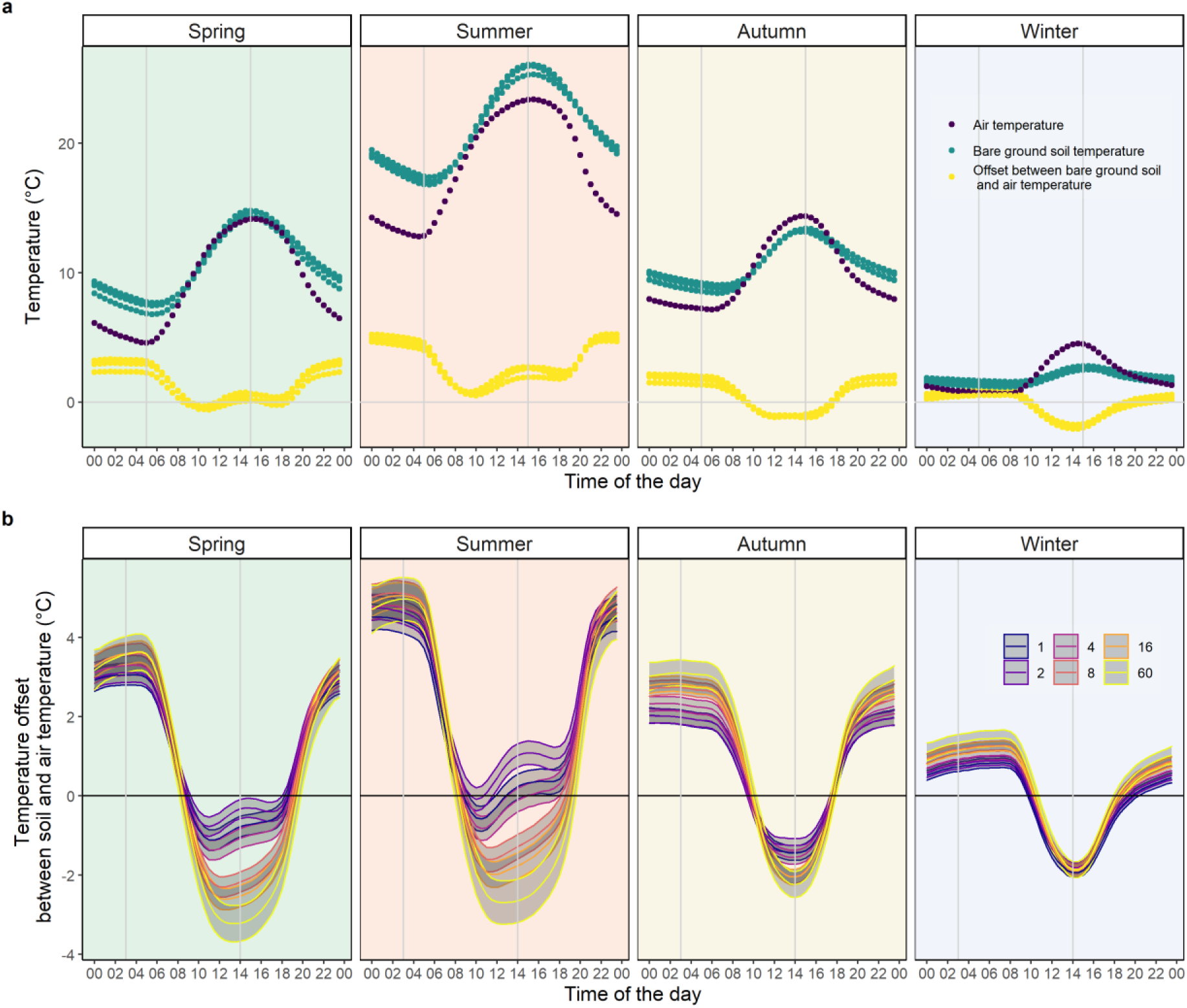
Daily temporal pattern of temperature offset between vegetated plot soil temperature and air temperature changes with plant diversity and season. **a,** The offset between the soil temperature of the four bare ground plots and the air temperature. **b,** The offset between the soil temperature of different vegetated plots with a gradient of plant diversity and air temperature.

**Extended Data Figure 4.**
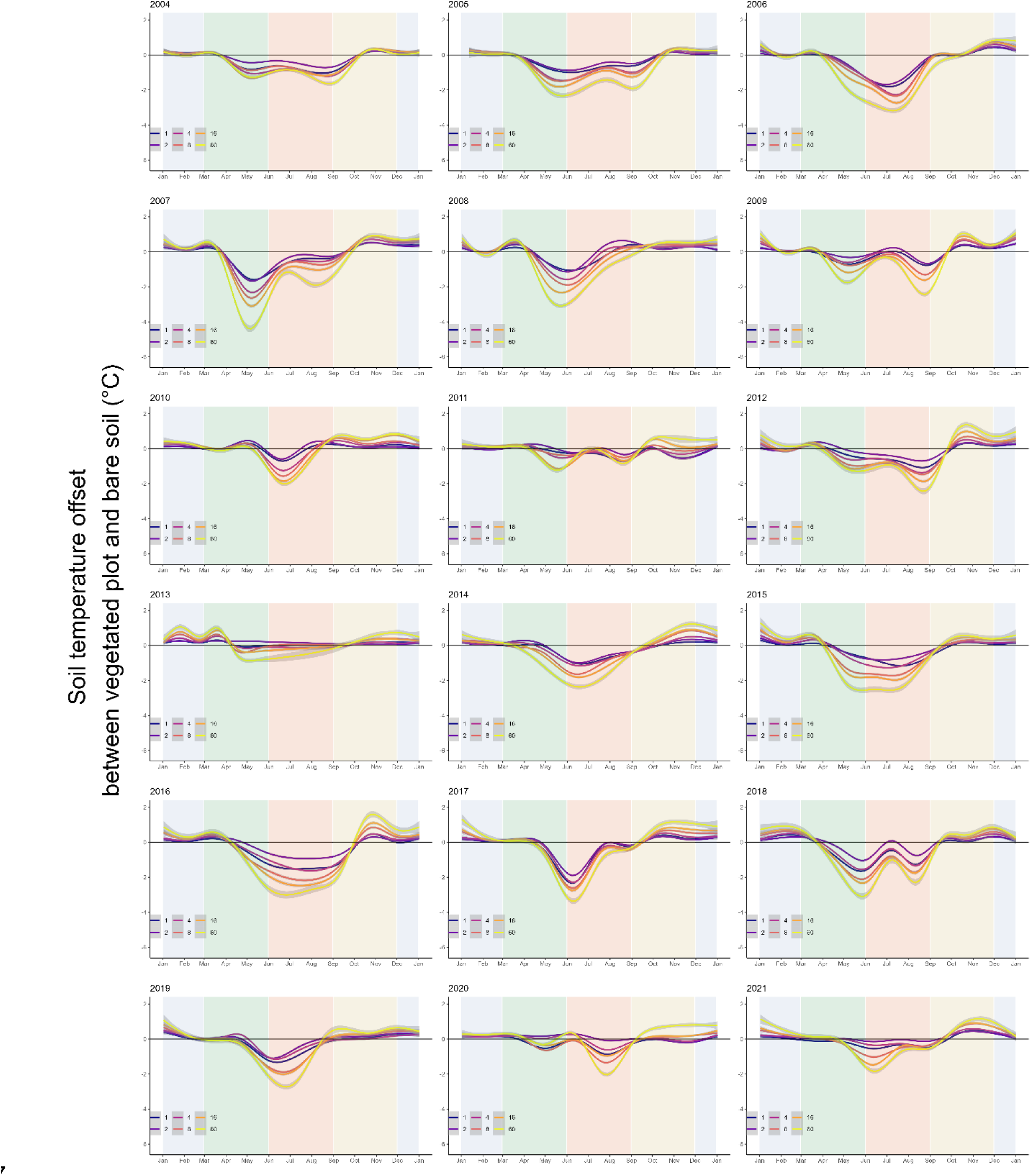
Offset of soil temperature at 5 cm between vegetated plots and bare soil at different plant diversity (1, 2, 4, 8, 16, and 60 species) on the daily scale for 18 years. Note that in 2013, the summer data (June, July and August) are missing due to the flood. So, the smoothing lines from June to August are not well represented in 2013.

**Extended Data Figure 4 5.**
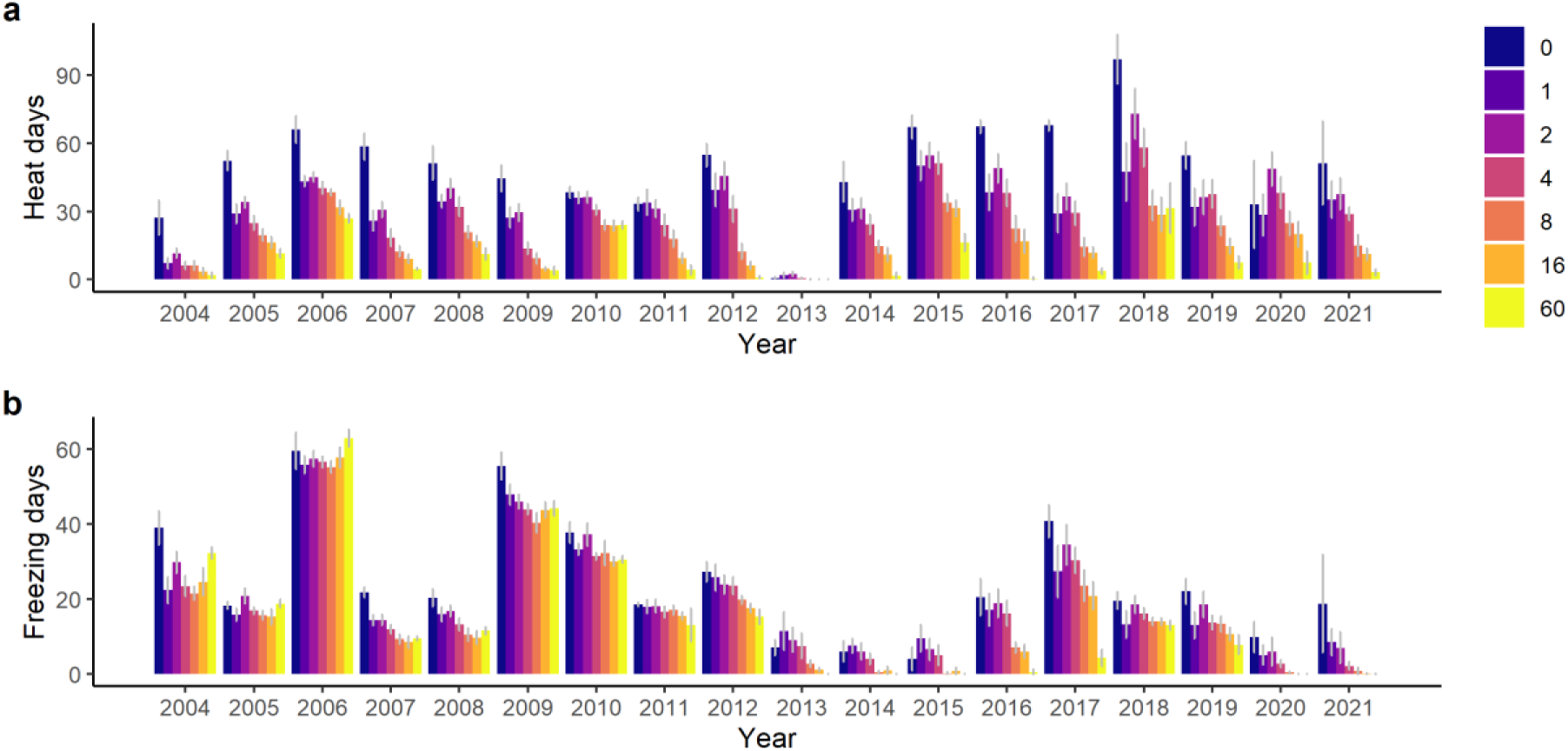
Extreme climate days in plant communities with different plant diversity levels in each year. The mean number of freezing days (minimum soil temperature at 5 cm is lower than 0°C) and standard error are shown in figure **a**. The mean number of heat days (maximum soil temperature at 5 cm is higher than 25°C) and standard error are shown in figure **b**. Note that in 2013, summer data (June, July and August) are missing due to the flood.

**Extended Data Figure 6.**
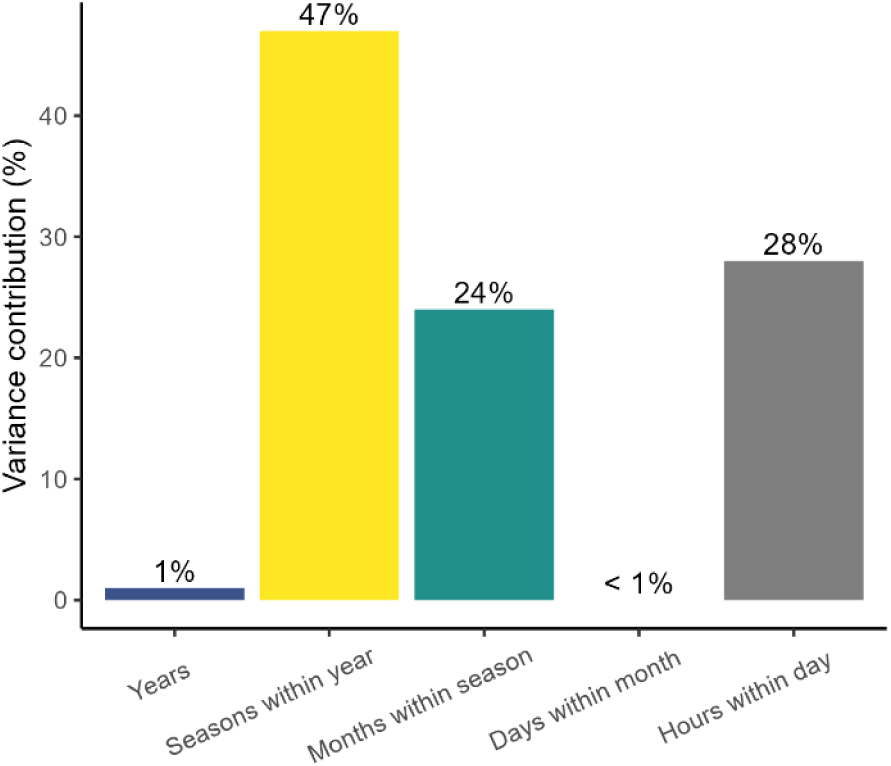
The hourly effect of plant diversity was calculated (24 hours per day, 365/366 days per year, 18 years, n = 157,800). After considering the effects of air temperature, the residual variance of the effects of plant diversity is decomposed into parts explained by different time scales.

**Extended Data Figure 7.**
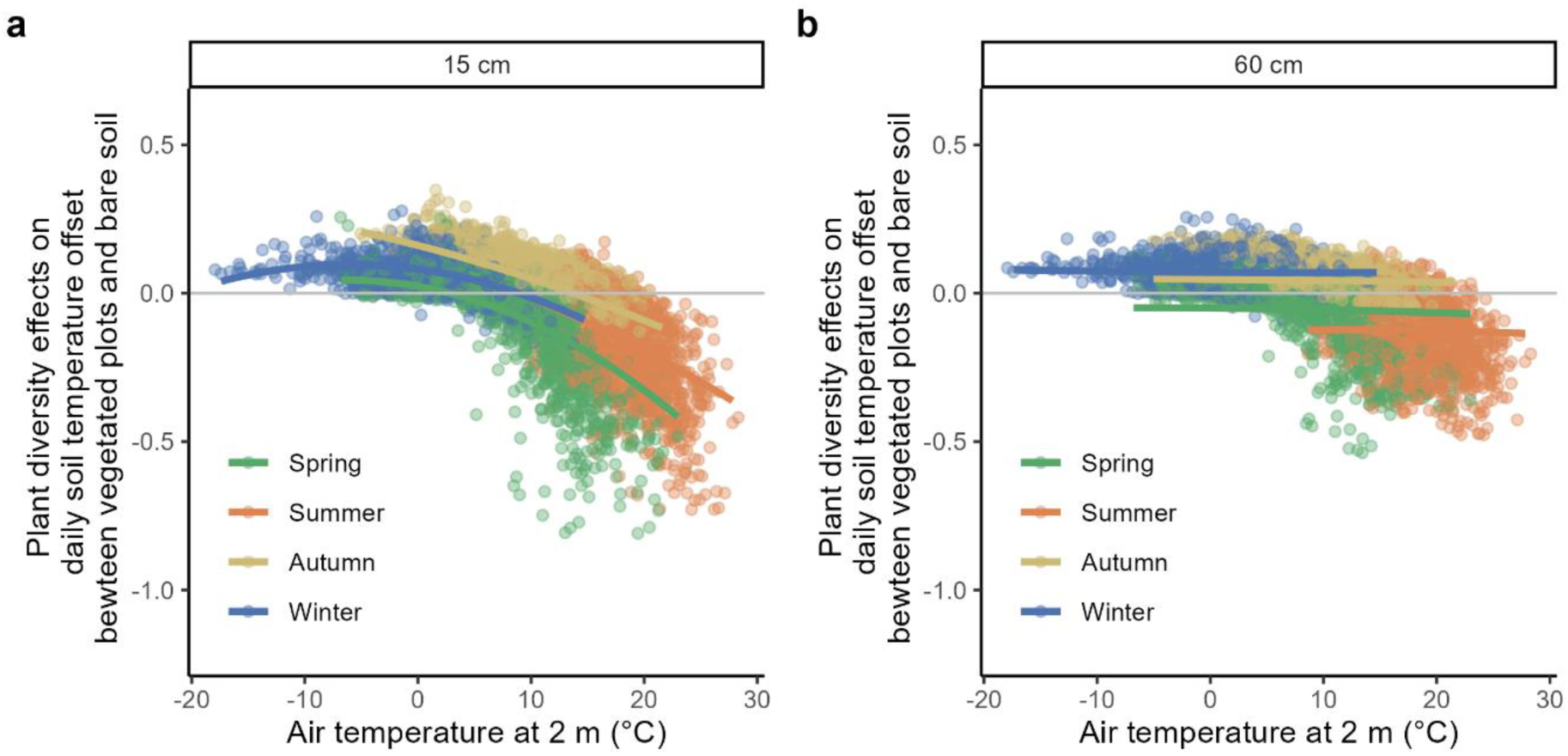
Relationship between air temperature and the effects of plant diversity at 15 cm (**a**) and 60 cm (**b**) soil depths. Solid lines are predicted data from the mixed-effects model.

**Extended Data Figure 8.**
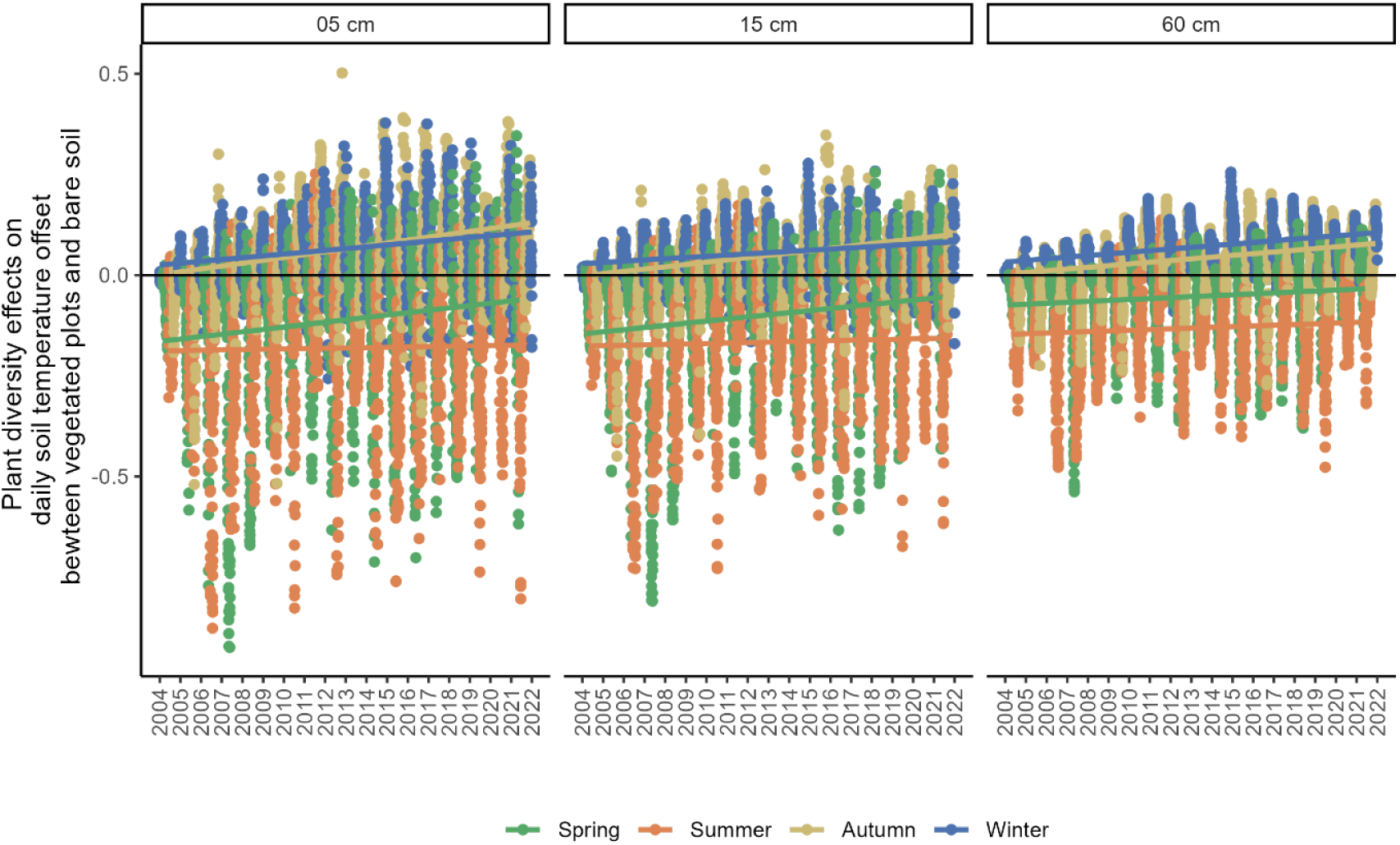
Effects of plant diversity change over time at different soil depths. The y-axis is the plant diversity effect on the differences between soil temperature in vegetation plots and bare ground. Solid lines are the effect trends for different seasons over time.

**Extended Data Figure 9.**
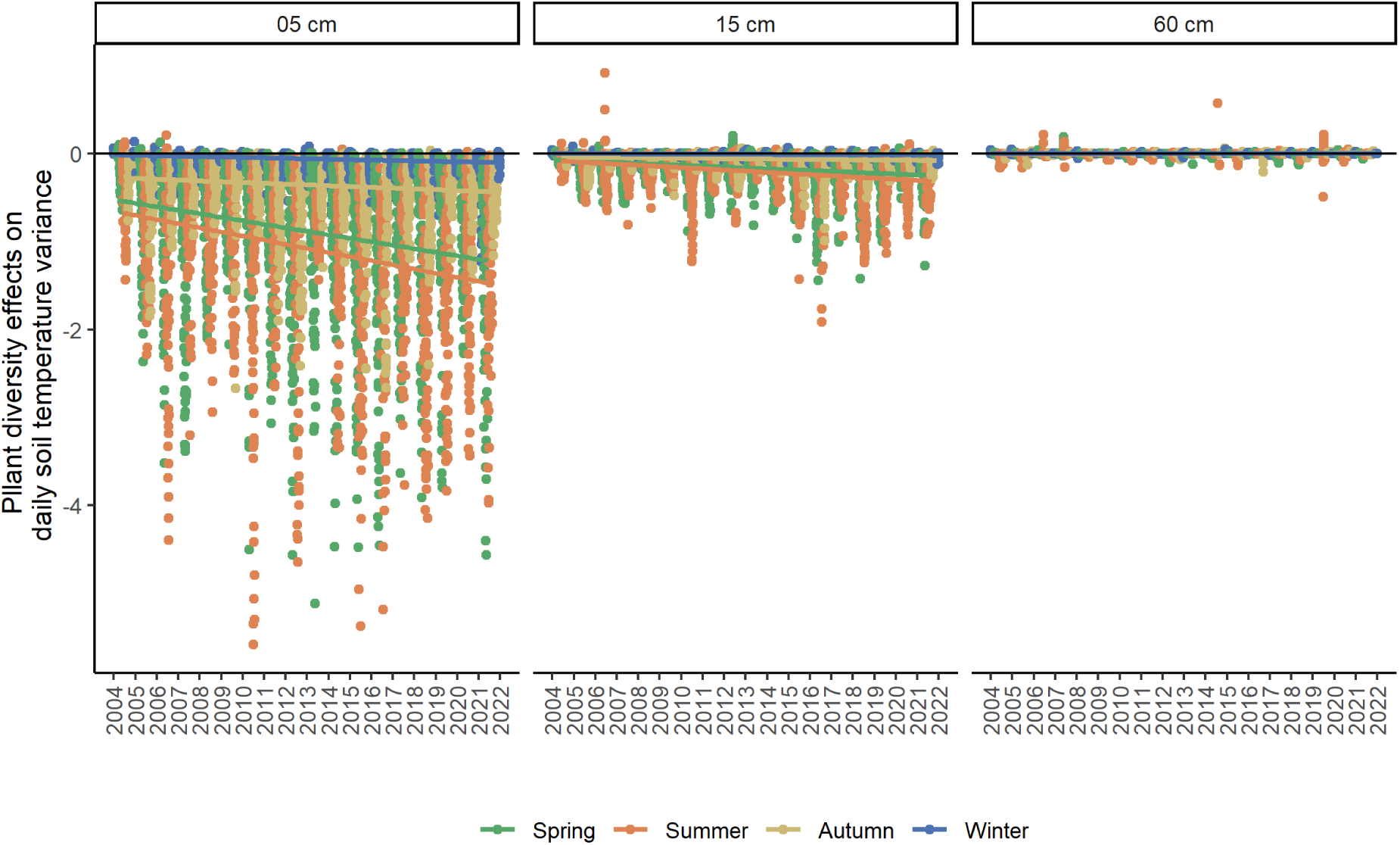
Plant diversity effects on the daily soil temperature variance change with time at different soil depths. Lines are mixed-effects model fits, with each color representing each season.

**Extended Data Figure 10.**
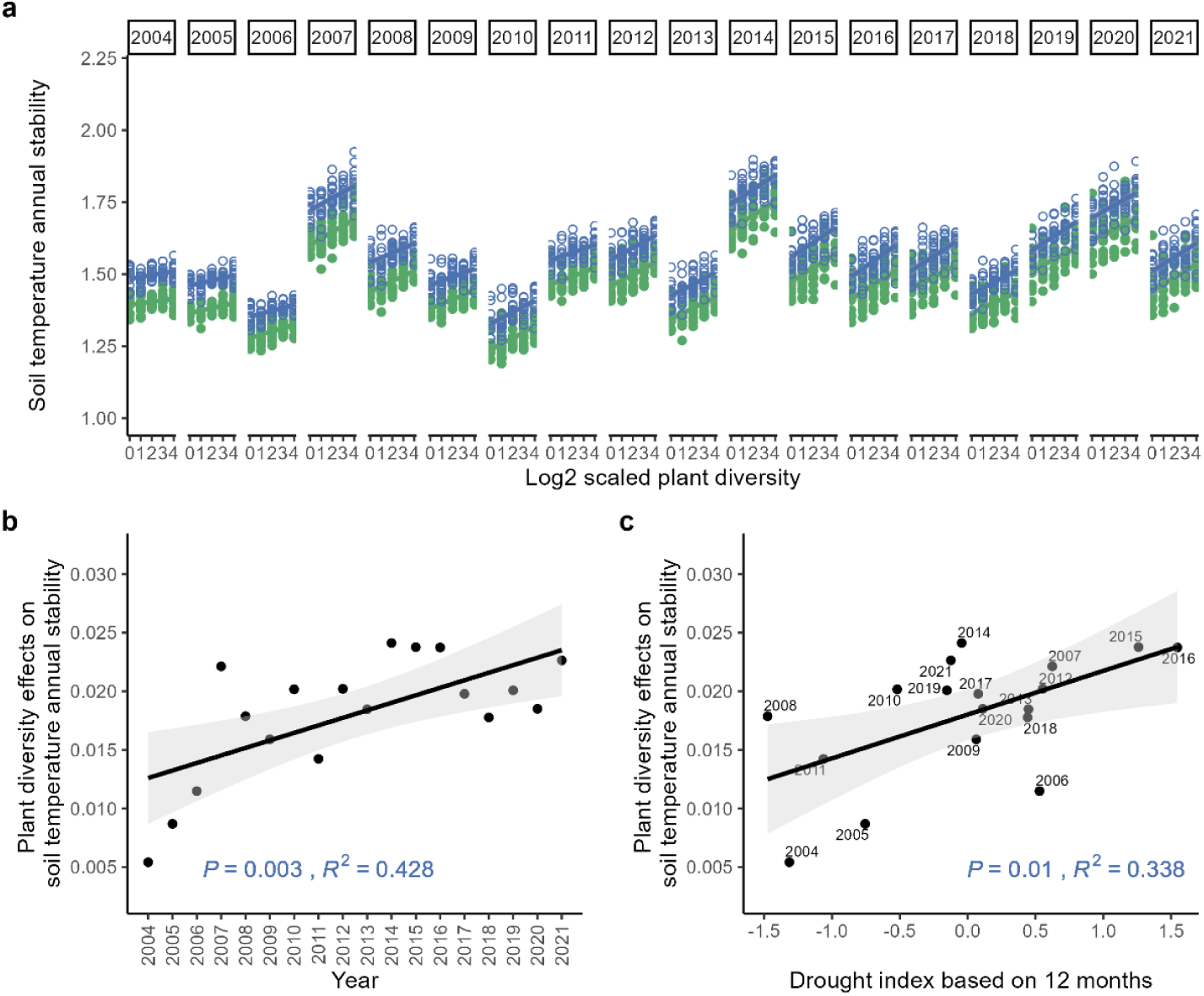
Effects of plant diversity on intra-annual soil temperature stability (**a**), and those effects change with time (**b**) and drought index (**c**). The 60-species diversity level data were excluded from this sensitivity analysis. The drought index here is calculated by multiplying the SPEI by -1, i.e. the drought situation becomes more severe with increasing values.

**Extended Data Figure 11.**
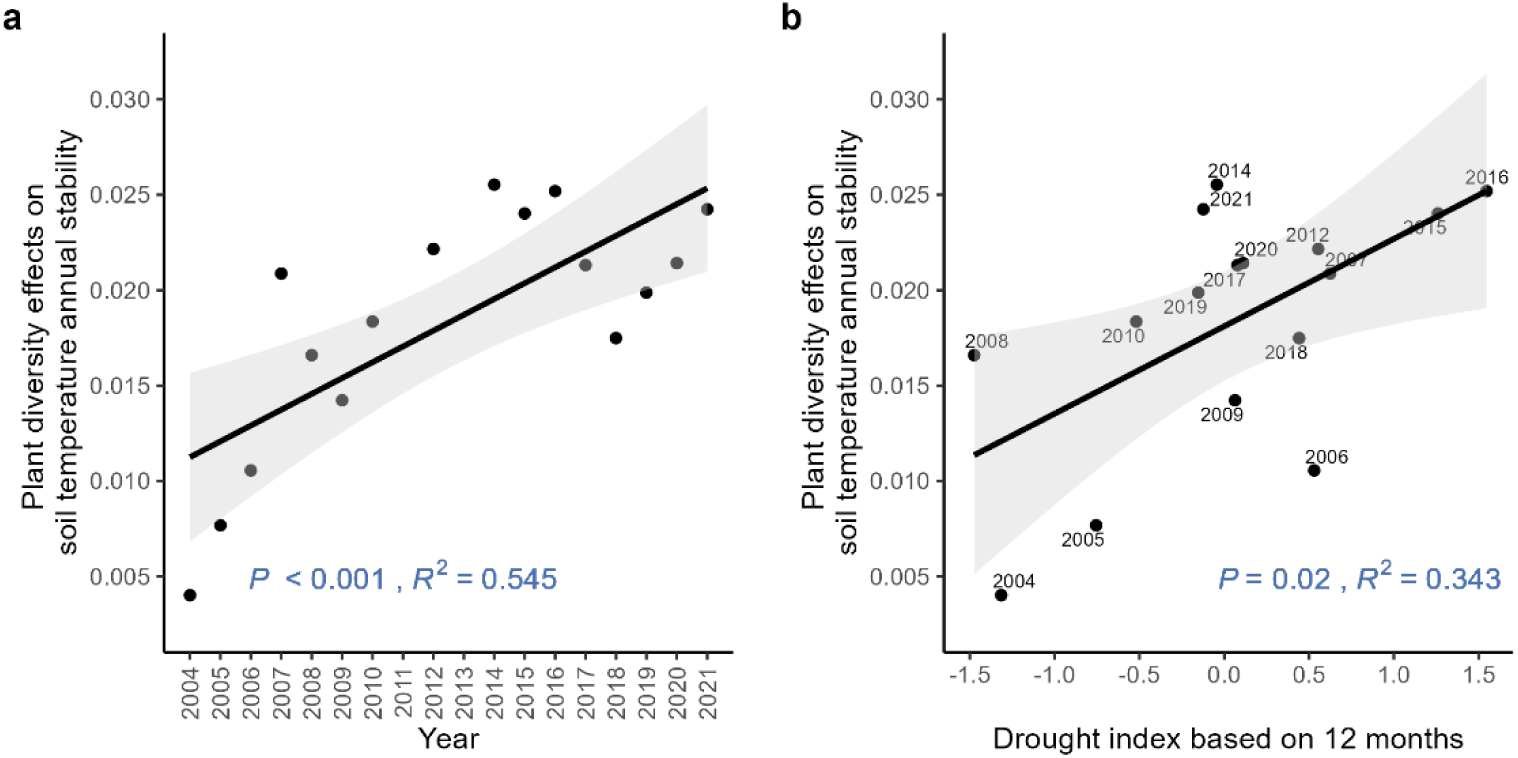
Effects of plant diversity change with time (**a**) and drought index (**b**). Here, we excluded those two years’ data (the year 2013 and 2011) that have high missing values. The drought index here is calculated by multiplying the SPEI by -1, i.e. the drought situation becomes more severe with increasing values.

**Extended Data Figure 12.**
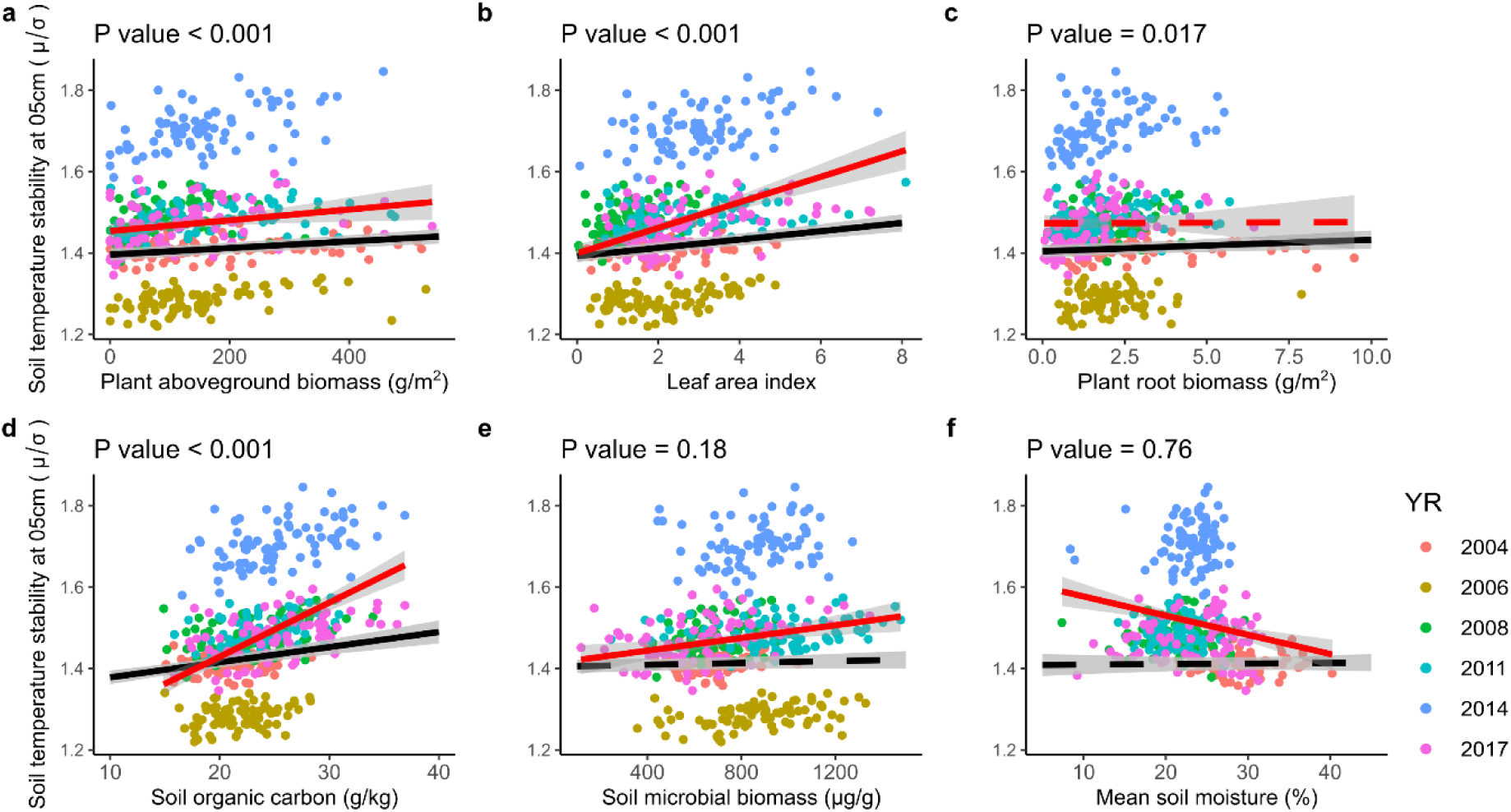
Relationships between different covariates and intra-annual soil temperature stability. Each closed circle represents one measurement, with different colours representing different years. The red line is the simple linear regression line of the selected variable and soil temperature stability (n = 480). In contrast, the black line is predicted from the mixed effects model after considering the effect of block and year. At the same time, the plot is also considered in the random term. The P values in the panels show the significance of the main effect of the variable on the x-axis from the mixed-effects model. Dashed lines indicate that the effect is not statistically significant, while solid lines represent significant effects.

**Extended Data Table 1.**
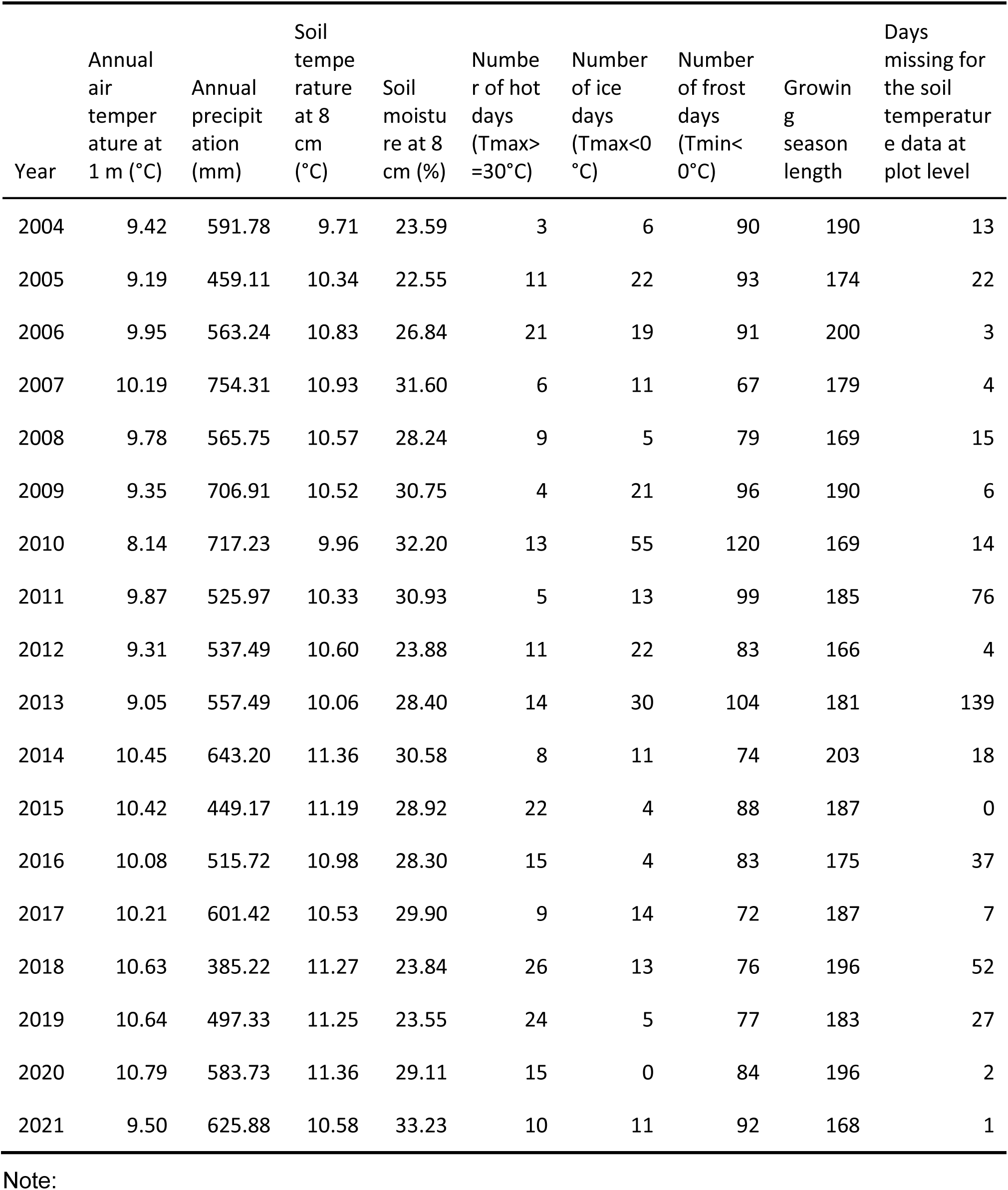
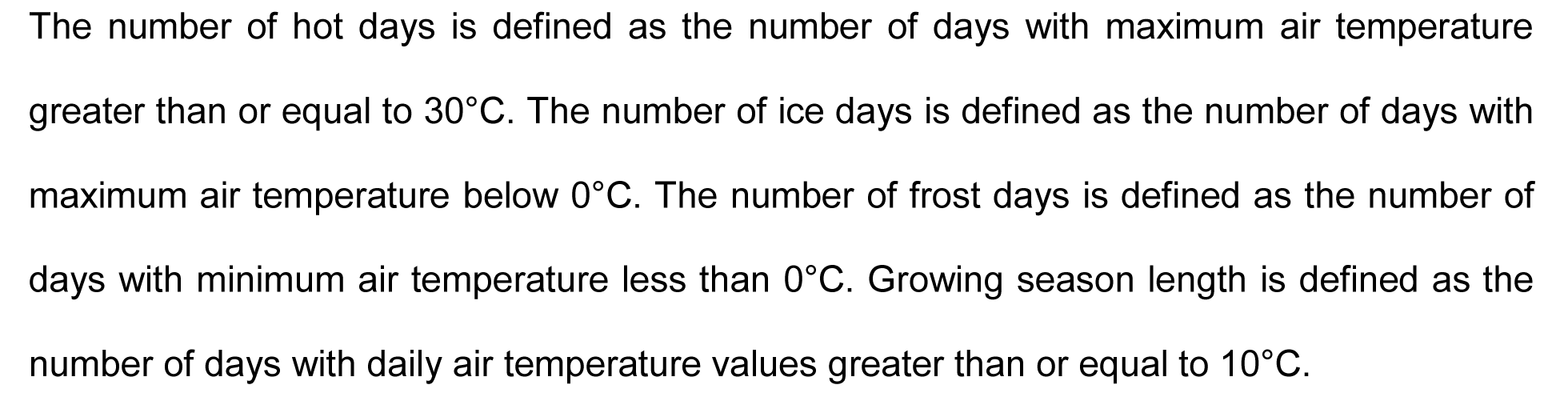
Summary of annual climate data and number of missing days for the soil temperature dataset per year.

| Year | Annual<br>air<br>temper<br>ature at<br>1 m (°C) | Annual<br>precipit<br>ation<br>(mm) | Soil<br>tempe<br>rature<br>at 8<br>cm<br>(°C) | Soil<br>moistu<br>re at 8<br>cm (%) | Numbe<br>r of hot<br>days<br>(Tmax><br>=30°C) | Number<br>of ice<br>days<br>(Tmax<0<br>°C) | Number<br>of frost<br>days<br>(Tmin<<br>0°C) | Growin<br>g<br>season<br>length | Days<br>missing for<br>the soil<br>temperatur<br>e data at<br>plot level |
| --- | --- | --- | --- | --- | --- | --- | --- | --- | --- |
| 2004 | 9.42 | 591.78 | 9.71 | 23.59 | 3 | 6 | 90 | 190 | 13 |
| 2005 | 9.19 | 459.11 | 10.34 | 22.55 | 11 | 22 | 93 | 174 | 22 |
| 2006 | 9.95 | 563.24 | 10.83 | 26.84 | 21 | 19 | 91 | 200 | 3 |
| 2007 | 10.19 | 754.31 | 10.93 | 31.60 | 6 | 11 | 67 | 179 | 4 |
| 2008 | 9.78 | 565.75 | 10.57 | 28.24 | 9 | 5 | 79 | 169 | 15 |
| 2009 | 9.35 | 706.91 | 10.52 | 30.75 | 4 | 21 | 96 | 190 | 6 |
| 2010 | 8.14 | 717.23 | 9.96 | 32.20 | 13 | 55 | 120 | 169 | 14 |
| 2011 | 9.87 | 525.97 | 10.33 | 30.93 | 5 | 13 | 99 | 185 | 76 |
| 2012 | 9.31 | 537.49 | 10.60 | 23.88 | 11 | 22 | 83 | 166 | 4 |
| 2013 | 9.05 | 557.49 | 10.06 | 28.40 | 14 | 30 | 104 | 181 | 139 |
| 2014 | 10.45 | 643.20 | 11.36 | 30.58 | 8 | 11 | 74 | 203 | 18 |
| 2015 | 10.42 | 449.17 | 11.19 | 28.92 | 22 | 4 | 88 | 187 | 0 |
| 2016 | 10.08 | 515.72 | 10.98 | 28.30 | 15 | 4 | 83 | 175 | 37 |
| 2017 | 10.21 | 601.42 | 10.53 | 29.90 | 9 | 14 | 72 | 187 | 7 |
| 2018 | 10.63 | 385.22 | 11.27 | 23.84 | 26 | 13 | 76 | 196 | 52 |
| 2019 | 10.64 | 497.33 | 11.25 | 23.55 | 24 | 5 | 77 | 183 | 27 |
| 2020 | 10.79 | 583.73 | 11.36 | 29.11 | 15 | 0 | 84 | 196 | 2 |
| 2021 | 9.50 | 625.88 | 10.58 | 33.23 | 10 | 11 | 92 | 168 | 1 |

Note:
The number of hot days is defined as the number of days with maximum air temperature greater than or equal to 30°C. The number of ice days is defined as the number of days with maximum air temperature below 0°C. The number of frost days is defined as the number of days with minimum air temperature less than 0°C. Growing season length is defined as the number of days with daily air temperature values greater than or equal to 10°C.

**Extended Data Table 2.**
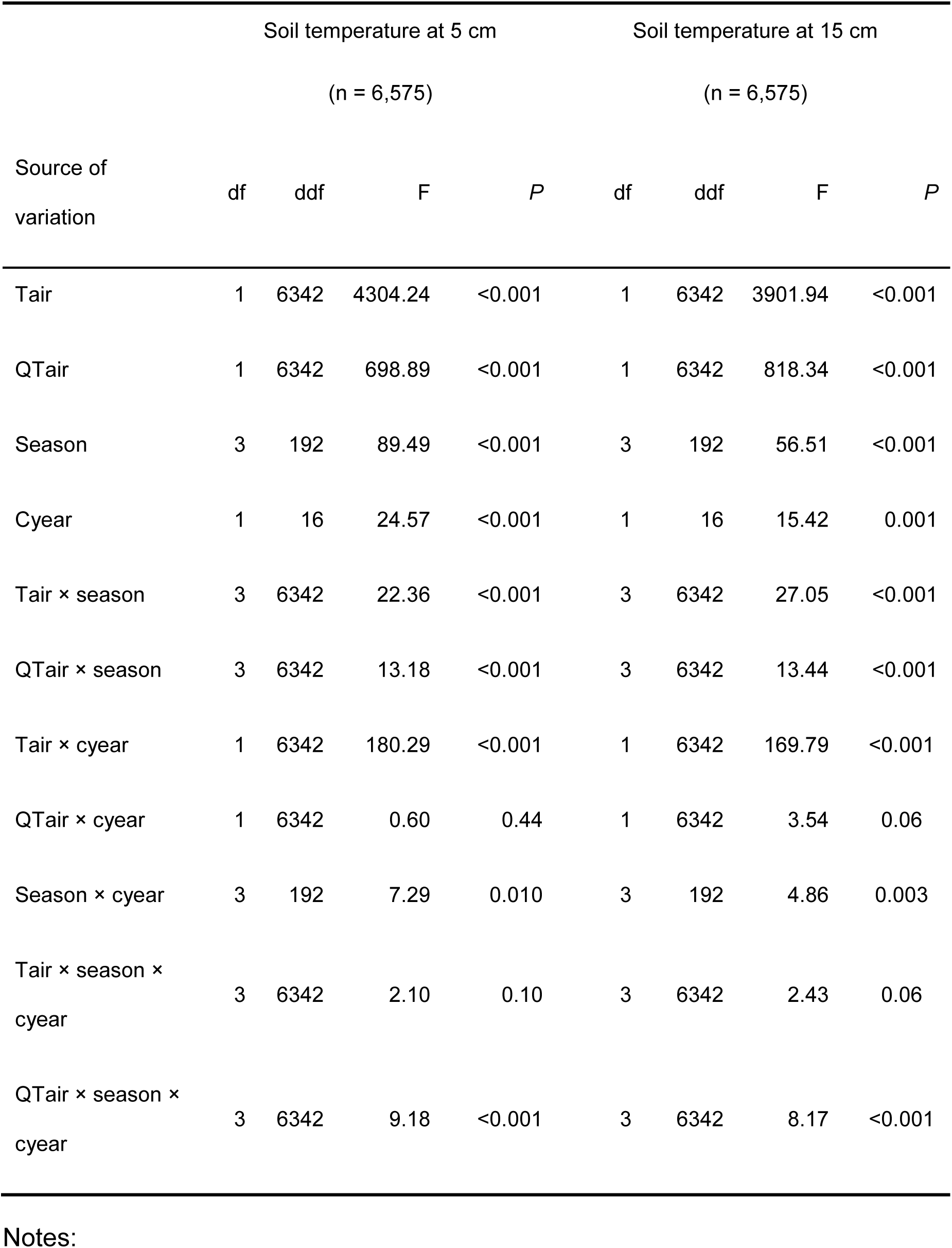
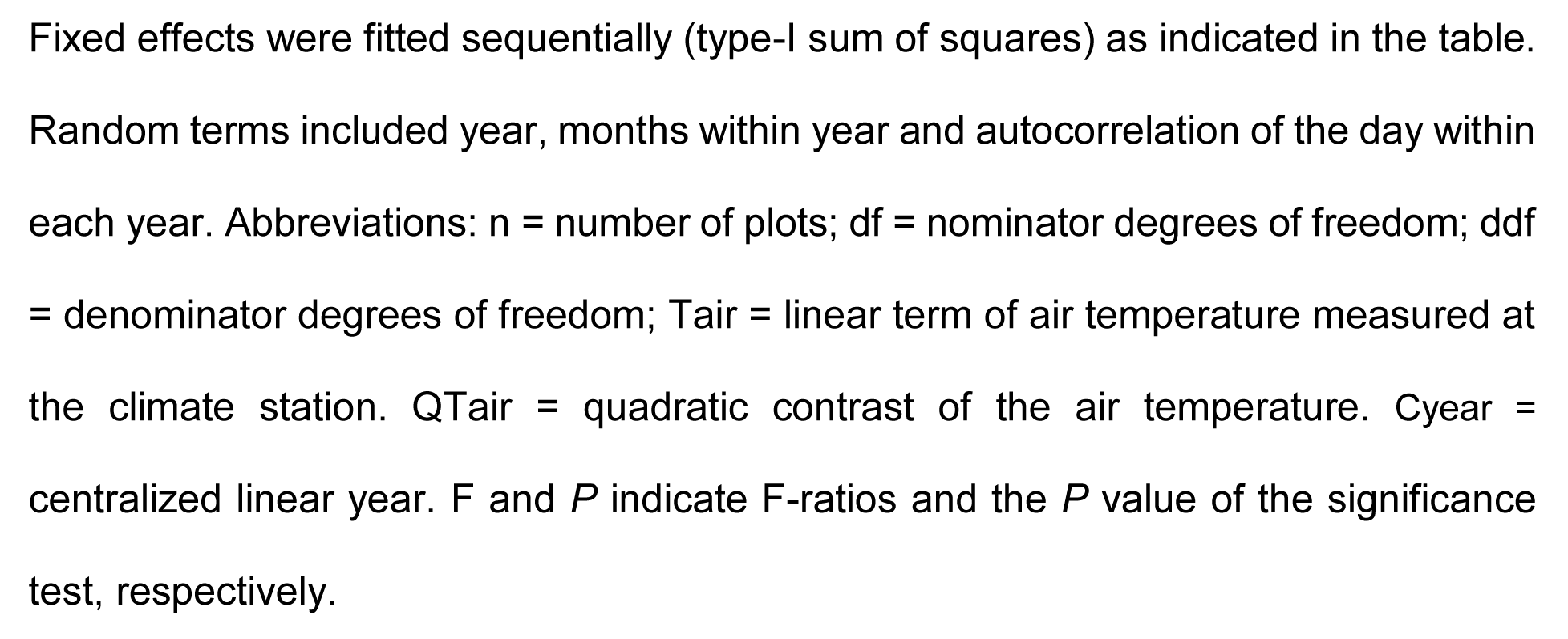
Mixed-effects models for the effects of air temperature, season, and year on the buffering effects of plant diversity on the soil temperature offset between vegetated and bare plots.

| Source of variation | Soil temperature at 5 cm |  |  |  | Soil temperature at 15 cm |  |  |  |
| --- | --- | --- | --- | --- | --- | --- | --- | --- |
|  | (n = 6,575) |  |  |  | (n = 6,575) |  |  |  |
|  | df | ddf | F | P | df | ddf | F | P |
| Tair | 1 | 6342 | 4304.24 | <0.001 | 1 | 6342 | 3901.94 | <0.001 |
| QTair | 1 | 6342 | 698.89 | <0.001 | 1 | 6342 | 818.34 | <0.001 |
| Season | 3 | 192 | 89.49 | <0.001 | 3 | 192 | 56.51 | <0.001 |
| Cyear | 1 | 16 | 24.57 | <0.001 | 1 | 16 | 15.42 | 0.001 |
| Tair × season | 3 | 6342 | 22.36 | <0.001 | 3 | 6342 | 27.05 | <0.001 |
| QTair × season | 3 | 6342 | 13.18 | <0.001 | 3 | 6342 | 13.44 | <0.001 |
| Tair × cyear | 1 | 6342 | 180.29 | <0.001 | 1 | 6342 | 169.79 | <0.001 |
| QTair × cyear | 1 | 6342 | 0.60 | 0.44 | 1 | 6342 | 3.54 | 0.06 |
| Season × cyear | 3 | 192 | 7.29 | 0.010 | 3 | 192 | 4.86 | 0.003 |
| Tair × season × cyear | 3 | 6342 | 2.10 | 0.10 | 3 | 6342 | 2.43 | 0.06 |
| QTair × season × cyear | 3 | 6342 | 9.18 | <0.001 | 3 | 6342 | 8.17 | <0.001 |

Notes:
Fixed effects were fitted sequentially (type-I sum of squares) as indicated in the table. Random terms included year, months within year and autocorrelation of the day within each year. Abbreviations: $n$ = number of plots; $df$ = nominator degrees of freedom; $ddf$ = denominator degrees of freedom; $T_{air}$ = linear term of air temperature measured at the climate station. $QT_{air}$ = quadratic contrast of the air temperature. $C_{year}$ = centralized linear year. $F$ and $P$ indicate F-ratios and the $P$ value of the significance test, respectively.

## Notes

### Competing Interest Statement

The authors have declared no competing interest.

